# Columnar cholinergic neurotransmission onto T5 cells of *Drosophila*

**DOI:** 10.1101/2024.08.28.610055

**Authors:** Eleni Samara, Tabea Schilling, Inês M. A. Ribeiro, Juergen Haag, Maria-Bianca Leonte, Alexander Borst

**Author notes:** Correspondence (E.S.), (A.B.).

## Abstract

Several nicotinic and muscarinic acetylcholine receptors (AChRs) are expressed in the brain of *Drosophila melanogaster*. However, the contribution of different AChRs to visual information processing remains poorly understood. T5 cells are the primary motion-sensing neurons in the OFF pathway and receive input from four different columnar cholinergic neurons, Tm1, Tm2, Tm4 and Tm9. We reasoned that different AChRs in T5 postsynaptic sites might contribute to direction selectivity, a central feature of motion detection. We show that the nicotinic nAChRα1, nAChRα4, nAChRα5 and nAChRα7 subunits localize on T5 dendrites. By targeting synaptic markers specifically to each cholinergic input neuron, we find a prevalence of the nAChRα5 in Tm1-, Tm2- and Tm4-to-T5 synapses and of nAChRα7 in Tm9-to-T5 synapses. Knock-down of nAChRα4, nAChRα5, nAChRα7, or mAChR-B individually in T5 cells alters the optomotor response and reduces T5 directional selectivity. Our findings indicate a differential contribution of postsynaptic receptors to input visual processing and, thus, to the computation of motion direction in T5 cells.

## Introduction

Neurotransmission is a key process of neuronal communication that entails the release of neurotransmitter from the presynaptic terminal and its binding by receptors on the postsynaptic membrane. In *Drosophila melanogaster*, the neurotransmitters acetylcholine, GABA, glutamate, dopamine, serotonin, octopamine^1^ and histamine^2^ are used for various neuronal tasks. Amongst them, the prime excitatory neurotransmitter is acetylcholine^3–5^, which can bind to a large number of structurally diverse acetylcholine receptors (AChRs)^6–9^. How this structural variety supports specific neural computations is not understood.

AChRs comprise two categories in bilaterians and cnidarians^10^: the fast ionotropic nicotinic receptors (nAChRs) and the slow metabotropic muscarinic receptors (mAChRs). nAChRs are activated by acetylcholine (ACh) and the agonist nicotine and belong to the cys-loop ligand-gated ion channel superfamily, forming functional pentamers of various subunit stoichiometries, homomeric or heteromeric. In *Drosophila* the pentameric nAChR formation is achieved from a pool of seven α and three β subunits^11–13^. mAChRs are G protein-coupled receptors (GPCRs), activated by ACh and the agonist muscarine and comprise three types, A, B and C in *Drosophila*^14–16^. These mAChR types activate different signaling cascades, with the mAChR-A and mAChR-C being coupled to the G_q/11_ signaling cascade and the mAChR-B to the G_i/0_^15,17,18^. Although the variety of AChRs has been thoroughly addressed at the biochemical level^18,19^, our understanding at the circuit level in terms of neural computations and their behavioral output is only starting to advance^14,15,20–23^.

Among sensory modalities, vision contributes the most synaptic input to the *Drosophila* central brain^24^. Visual processing starts at the two compound eyes, whose retinotopic organization leads to a repetitive circuitry across its approximately 850 facets^25,26^. The major neuroanatomical area dedicated to visual information processing is the optic lobe, which is divided in four neuropils, the lamina, the medulla, the lobula and the lobula plate. From all the different visual cues such as color and polarization vision^27^, motion vision has been the one thoroughly investigated, mainly due to its importance in courtship and navigation of the individual^28–30^. The photoreceptor-derived information is split into two pathways: the ON-pathway for luminance increments and the OFF-pathway for luminance decrements^31,32^. Along the motion vision circuitry, the first neurons that compute directional information are T4 cells in the ON-pathway and T5 cells in the OFF-pathway^33,34^.

Both T4 and T5 cells share the same bifurcation morphology, with the dendrite giving rise to the axonal terminal fiber and to the soma fiber^35^. The T4 dendrites are located in medulla layer ten (M10) and the T5 dendrites in lobula layer one (LO1). T4 and T5 cells exist in four subtypes per column: a, b, c and d. Each subtype has its dendrite oriented in one of the four cardinal directions and responds selectively to front-to-back, back-to-front, upward or downward motion, respectively^34–37^. Furthermore, each subtype has its axonal terminals in one of the four layers in lobula plate^35^. For their directional responses, T4 and T5 cells rely on the visual information they receive from input neurons. The major upstream columnar neurons of T4 cells are the excitatory cholinergic Mi1 and Tm3 neurons, the glutamatergic Mi9 and the inhibitory GABAergic CT1, C3 and Mi4 neurons^38–46^. For T5 cells, the main columnar input neurons are the excitatory cholinergic Tm1, Tm2, Tm4 and Tm9 cells and the inhibitory GABAergic CT1 neuron^40,42–44,47–52^. T4 and T5 cells collect their inputs in a distinct spatial order^40^ reflecting their specific directional tuning^53,54^.

T4 and T5 cells achieve their response selectivity for a particular direction by a combination of two mechanisms: preferred direction enhancement and null direction suppression^54,55^. In both processes, the luminance signal from one point in visual space is delayed and interacts with the instantaneous signal from a neighboring image point. For preferred direction enhancement, the delayed signal is from the preferred side of the cell, i.e. where a stimulus moving in the preferred direction enters the receptive field of the cell, and amplifies the central signal. For null direction suppression, the delayed signal is from the opposite, the so-called null side, and is used to suppress the central signal. In T4 cells, preferred direction enhancement is biophysically implemented by a release from shunting inhibition, mediated by the glutamatergic, inhibitory OFF-center cell Mi9, which amplifies the central excitatory input from Mi1 and Tm3 neurons^45,56^. For null direction suppression, the inhibitory neurons CT1, Mi4 and C3 act in concert to inhibit the Mi1-Tm3 signal. In T5 cells, the CT1 cell is responsible for null direction suppression^51^. It is, however, unclear how the different cholinergic inputs accomplish a preferred direction enhancement. While the functional consequences of the different T5 cholinergic input neurons have been addressed in the past^48,52,57^, the functional implications of a possible input-specific AChR composition on T5 dendrites have not been considered.

In this study, we address the structural and functional variety of different AChRs in T5 cells. We first show the existence of the nAChRα1, nAChRα4, nAChRα5 and nAChRα7 subunits on T5 dendrites. We target synaptic markers to each T5 columnar cholinergic input and find a differential distribution of AChRs in T5 postsynaptic sites. Through a combinatorial approach of neuronal input spatial wiring and synaptic receptor mapping, we demonstrate the complex columnar cholinergic spatial wiring on T5 dendrites. Finally, we show that the nAChRα4, nAChRα5 and nAChRα7 subunits, as well as the mAChR-B type, differentially affect the optomotor response and the directional responses of T5 cells.

## Results

### Expression of AChRs in T5 cells

Information about the expression of AChRs in T5 cells is available at the RNA level^43,58,59^, but is lacking at the protein level. We first evaluated the endogenous expression of the nAChRα1, nAChRα2, nAChRα4, nAChRα5, nAChRα6 and nAChRα7 subunits on T5 dendrites (Figure 1A,B). We used nAChRα subunits endogenously tagged with an enhanced version of the green fluorescent protein (EGFP) generated by Pribbenow et al.^22^, thus minimizing the mistargeting error that can occur with receptor overexpression and allowing for the assessment of protein expression levels. We verified the localization of nAChRs in T5 neurons by co-localizing the receptor puncta with the membrane-bound tdTomato expressed in T5 dendrites. We found the nAChRα1, nAChRα4, nAChRα5, nAChRα6 and nAChRα7 subunits on T5 dendrites, while the nAChRα2 subunit was not detected (Figure 1C-I). To ensure that the lack of nAChRα2 expression was T5-specific, we confirmed the nAChRα2 expression in other LO layers in the optic lobe (Figure S1A). By quantifying the nAChRα subunit density in regions of interest (ROIs) on T5 dendrites in LO1, we observed a prevalence of the nAChRα5 subunit and low co-localization levels of the nAChRα2 and the nAChRα6 subunits with T5 dendrites, which is in agreement with previous nAChRα RNA expression levels^43,58^ (Figure 1J, Figure S1B-D). A recent study^60^ also confirmed the T5 dendritic localization of the nAChRα3 and nAChRβ1 subunits, which together with our nAChRα1, nAChRα4, nAChRα5, nAChRα6 and nAChRα7 subunit data, emphasizes the large variety of nAChRs present on T5 dendrites. Next, we sought to identify the nAChRα expression patterns in the T5 axonal terminals by searching for the co-localization of receptor puncta with the membrane-bound tdTomato expressed in T5 terminals. Both T4 and T5 neuronal populations were targeted. Due to the lack of neuropile distinction between T4 and T5 axonal terminals, our results cannot be attributed explicitly to T5 cells. The nAChRα1, nAChRα5, nAChRα6 and nAChRα7 subunits were detected in T4-T5 axonal terminals, while the nAChRα2 and nAChRα4 were not (Figure S2A-G). Additionally, we observed the reduction of the nAChRα1, nAChRα5 and nAChRα7 subunits in LOP after a T4-T5 specific subunit knock-down, indicative of their T4-T5 axonal synaptic localization (Figure S2J, S3A,C,D).

**Figure 1.**
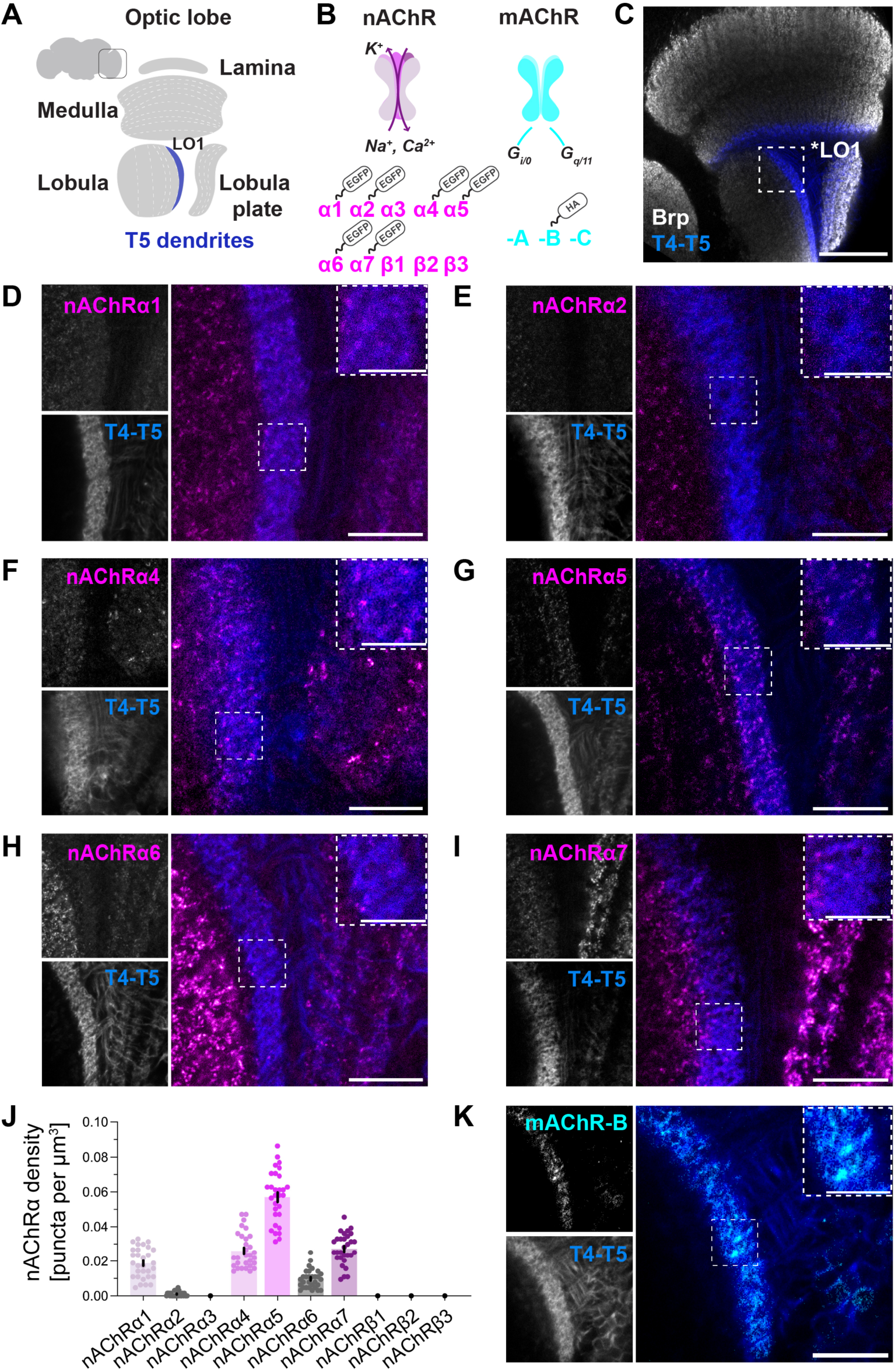
AChR expression on T5 dendrites. (A) Schematic representation of T5 dendrites residing in lobula layer 1 (LO1) of the *Drosophila* optic lobe (OL). (B) Schematic representation of nicotinic ionotropic (magenta) and muscarinic metabotropic AChRs (cyan) in *Drosophila*. Out of the seven nAChRα subunits, six were tagged with EGFP^22^. The mAChR-B was tagged with the HA peptide tag (C.H.Lee) (C) OL (grey) with T4 and T5 dendrites (R42F06-Gal4>UAS-myr::tdTomato, blue) in M10 and LO1 accordingly. Scale bar 40μm. (D-Ι) nAChRα1, nAChRα2, nAChRα4, nAChRα5, nAChRα6 and nAChRα7 (nAChRα::EGFP, magenta) subunit expression on T5 dendrites (R42F06-Gal4>UAS-myr::tdTomato, blue). Scale bar 10μm, inset 5μm. (J) Density of nAChRα subunit expression on T5 dendrites across 30 ROIs in LO1 (optic lobes n=6). nAChRα3, nAChRβ1, nAChRβ2 and nAChRβ3 subunits were not addressed. Subunits with the lowest expression levels are in grey. Data is mean ± SEM. (K) Overexpression of mAChR-B (UAS-mAChR-B::HA, cyan) on T5 dendrites (R42F06-Gal4>UAS-myr::tdTomato, blue). Scale bar 10μm, inset 5μm.

To study the mAChR expression on T5 dendrites, we used a line expressing the mAChR-B type under control of UAS. We observed the overexpressed mAChR-B protein localizing exclusively on T5 dendrites (Figure 1K, S2H). The encountered dendritic T5 mAChR-B type targeting might be the outcome of ectopic expression leading to dendritic protein targeting. We reasoned that if mAChR-B was mistargeted to T5 dendrites, this mistargeting would be observed in other cells and that over-expression of mAChR-B with a pan-neuronal driver would result in the presence of ectopic mAChR-B in all the dendrites of neurons in the fly brain (Figure S2I). Instead, the mAChR-B type followed an enrichment in layer 10, where T4 dendrites reside, and in LO1, where T5 dendrites reside, while being absent from other medulla layers and the lobula plate. In summary, the dendritic and axonal terminal protein expression patterns differ between ionotropic and metabotropic AChRs and even within the same AChR (Figure S2K).

To investigate the expression of those AChRs where we lacked endogenously tagged lines, and to verify the expression profiles reported in Figure 1, we used enhancer-trap Gal4 lines^61^ (Figure S4A-L). By comparing RNA sequencing datasets^43,58^ with the enhancer trap and the endogenously tagged experiments here and in previous studies^60^, we concluded that the nicotinic nAChRα1, nAChRα3, nAChRα4, nAChRα5, nAChRα7 and nAChRβ1 subunits as well as the muscarinic mAChR-A and mAChR-B types are expressed in T5 cells, while the nAChRα2, nAChRα6, nAChRβ2 and nAChRβ3 subunits exhibit low expression levels or are not expressed at all (Figure S4M). Transcriptomics suggest low mAChR-C expression levels in the fly brain^14,20^. Together our data show that the dendritic and axonal protein localization patterns of ionotropic and metabotropic AChRs in T5 cells are diverse and complex.

### Morphological identification of cholinergic T5 input neurons and synapse visualization

The nAChRα1, nAChRα4, nAChRα5 and nAChRα7 subunits might contribute to the directional responses of T5 subtypes by localizing to all cholinergic synapses linking Tm1, Tm2, Tm4 and Tm9 to T5 dendrites, or to only a subset of these. To discern between these possibilities, we assessed the protein localization of nAChRα subunits in specific synaptic pairs. We first needed genetic access to the four T5 cholinergic neuronal inputs: Tm1, Tm2, Tm4 and Tm9. The morphology of these input neurons has been extensively studied via Golgi staining^35^ and electron microscopy reconstructions^39,40,47,50,62–64^. We used this previously attained EM morphological knowledge to cross-validate our light microscopy screening of potential Tm-specific enhancer lines (Figure 2A,D,G,J). Both Gal4 and LexA driver lines^65,66^, previously used in functional studies^48,51^, were tested and their efficacy was assessed in terms of neuronal specificity and whole population targeting. Neuronal specificity was evaluated based on the arborization pattern of each neuronal type and the lobula layer where the axonal terminals are localized. To achieve sparse and stochastic neuronal labelling, we used the Multi-color FlpOut approach^67^ and observed axonal terminals in lobula layer 1 for Tm1 (Figure 2A), up to layer 2 for Tm2 (Figure 2D), up to layer 4 for Tm4 (Figure 2G), and in layer 1 for Tm9 (Figure 2J), as expected from previous studies^35,40^. To exclude the developmental bias as well as the non-neuronal targeting induced by the heat-shock-activated flipase^68^ in the Multi-color FlpOut approach (Figure 2G), we supplemented our single cell labelling with whole population labelling of the Tm-Gal4 lines (Figure 2B,E,H,K). This approach revealed the Gal4 driver lines expressing in a cell-type specific way in the whole population of Tm1, Tm2, Tm4 and Tm9 cells, respectively. For the respective LexA^66,69^ driver lines, the whole Tm1 population was successfully targeted, but after comparing the results with the analogous Gal4 driver line, dendritic ramifications in layer 4 that do not correspond to Tm1, were observed (Figure 2C). The Tm2, Tm4 and Tm9 LexA lines displayed sparse neuronal labelling, indicating that the driver lines could not target the whole neuronal population (Figure 2F,I,L). Based on this morphological screening, we therefore chose the Gal4 driver lines for the Tm-T5 synapse visualization.

**Figure 2.**
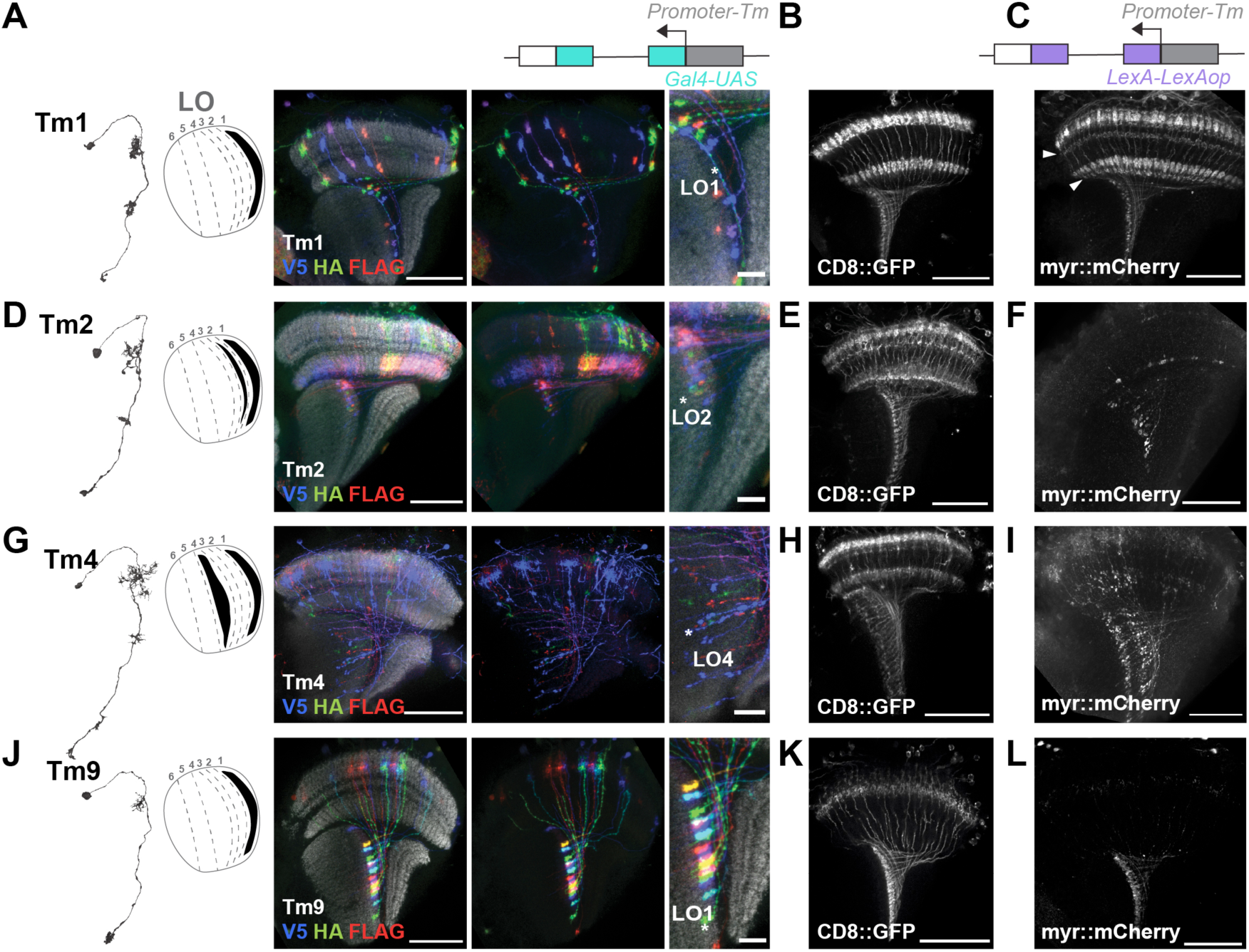
Morphological identification of the four columnar T5 cholinergic input neurons. (A) Electron microscopy (EM)(flywire.ai) reconstruction of a Tm1 neuron reveals two loci of arborization in medulla. Schematic representation of the Tm1 axonal terminal localization in LO1. Tm1-Gal4 expression (cyan) of the MultiColor FlpOut cassette (V5, HA, FLAG) and axonal localization in LO. Scale bar 40μm and 10μm. (B) Whole population labelling of Tm1 neurons (R74G01-Gal4>UAS-CD8::GFP). Scale bar 40μm. (C) Whole population labelling of Tm1-LexA (purple) neurons (VT041034-LexAGAD>LexAop-myr::mCherry). Arrows represent additional dendritic ramifications that lack in (B). Scale bar 40μm. (D-F) As in (A-C), but for Tm2 neurons (VT012282-Gal4, VT058650-LexAGAD). (G-I) As in (A-C), but for Tm4 neurons (R35H01-Gal4, R53C02-LexAGAD). (J-L) As in (A-C), but for Tm9 neurons (VT065303-Gal4, R24C08-LexAp65).

To visualize synapses between Tm1, Tm2, Tm4, Tm9 and T5 cells, we screened for the available reporter tools. First, we sought to explore methods of trans-synaptic labelling such as GFP Reconstitution Across Synaptic Partners (GRASP)^70^, where the GFP_1-10_ fragment is targeted to the cell membrane of the presynaptic compartment, while the GFP_11_ fragment is targeted to the cell membrane of the postsynaptic compartment. The targeted GRASP (t-GRASP)^71^ allowed to capture all synapses between Tm and T5 cells (Figure S5A). The presence of reconstituted GFP signal in the inner optic chiasm, likely due to the ectopic reconstitution in the Tm4-T5 synapses, led us to exclude this genetic method. The next reporter tool was that of the activity GRASP (syb-GRASP)^72^. The reconstituted GFP signal was indicative of the given synaptic activity at the point of the dissection, as the GFP_1-10_ fragment is tethered to the vesicular protein synaptobrevin, thus resulting in the syb-GRASP reconstitution being sparser than the t-GRASP reconstitution (Figure S5B). This approach did not allow for a quantitative analysis; therefore syb-GRASP was not chosen for our experiments. Finally, the truncated version of the presynaptic marker Brp, one of the major protein participants of the protein-dense T-bar^73,74^, named Brp^short^ ^75,76^ was also tested (Figure S5C). This Brp^short^::GFP line labelled only the presynaptic sites of all the Tm synapses, regardless of their activity, without labelling their postsynaptic sites. In conclusion, our reporter tool screening for synaptic labelling pointed to Brp^short^ as a suitable marker to evaluate the synaptic localization of the nAChRα receptor subunits.

To validate the synaptic specificity of Brp^short^, we assessed two different, well described synapses: the cholinergic Tm3-T4 and the glutamatergic Mi9-T4 synapse (Figure S6A,B). Brp^short::mCherry^ ^77^ successfully showed the expected lack of trans-synaptic alignment with the GABAergic receptor Rdl in Tm3-T4 synapses (Figure S6A), while it revealed trans-synaptic alignment with the glutamatergic receptor GluClα^45^ in Mi9-T4 synapses (Figure S6B). To confirm the synaptic localization of Brp^short::mStraw^, a chimera that has the same truncated form of Brp as in Brp^short::mCherry^ but is instead fused with mStrawberry by Owald et al.,^78^, we used an antibody against the endogenous Brp having affinity to amino acids absent from the truncated Brp^short^ form (Figure S6C). This approach revealed the pre-synaptic alignment of the endogenous Brp (Anti-Brp) with the Tm1, Tm2, Tm4 and Tm9 boutons (Brp^short::mStraw^). These results, together with previous studies^75^, speak in favor of Brp^short::mStraw^ as a valid pre-synaptic marker in our circuit of interest.

### The synaptic localization of the nAChR**α**5 subunit differs among the four cholinergic inputs

With Brp^short::mStraw^ as a presynaptic marker, we evaluated the localization of the most highly expressed subunits, nAChRα1, nAChRα4, nAChRα5 and nAChRα7, along the Tm-T5 synapses on T5 dendrites. We focused on the Tm1, Tm2, Tm4 and Tm9 synapses in LO1, the major cholinergic inputs to T5. The regions of interest were restricted to LO1 so as to include the Tm-T5 synaptic connections (Figure S7A). We found that the nAChRα1, nAChRα4, nAChRα5 and nAChRα7 subunits localize to Tm1, Tm2, Tm4 and Tm9 synapses in LO1 (Figure 3A-D, Figure S7B). The synapses of Tm cells in LO1 might account for other output neurons apart from T5 cells. To test this, we used the FlyWire connectome data explorer (Codex) for the full adult fly brain (FAFB) dataset^62–64^ in order to find the output neurons of the Tm1, Tm2, Tm4 and Tm9 synapses in LO1. This analysis revealed T5 cells as the principal output neuronal type across all four Tm cell types in LO1 (Figure S8A). Therefore, the quantification of the nAChRα subunit synaptic localization in LO1 principally corresponded to Tm-T5 synapses. Importantly, we found that not all Tm input neurons had the same synaptic receptor profiles. Tm1-, Tm2- and Tm4-to-T5 synapses primarily used the nAChRα5 subunit, whereas Tm9-to-T5 synapses mainly used the nAChRα7 subunit (Figure 3E-H). As the GABAergic neuron CT1 was a considerable Tm9 output in terms of synapse number in LO1 (Figure S8A), we wondered if the Tm9-nAChRα7 subunit trans-synaptic alignment (Figure 3H) could actually be attributed to Tm9-T5 synapses as well. We generated new flies to employ a C-RASP approach. C-RASP follows the activity GRASP principles, but instead of a GFP_1-10_ fragment uses a CFP_1-10_ fragment. This method showed the nAChRα7 subunit localization in Tm9-T5 synapses (Figure S8B).

**Figure 3.**
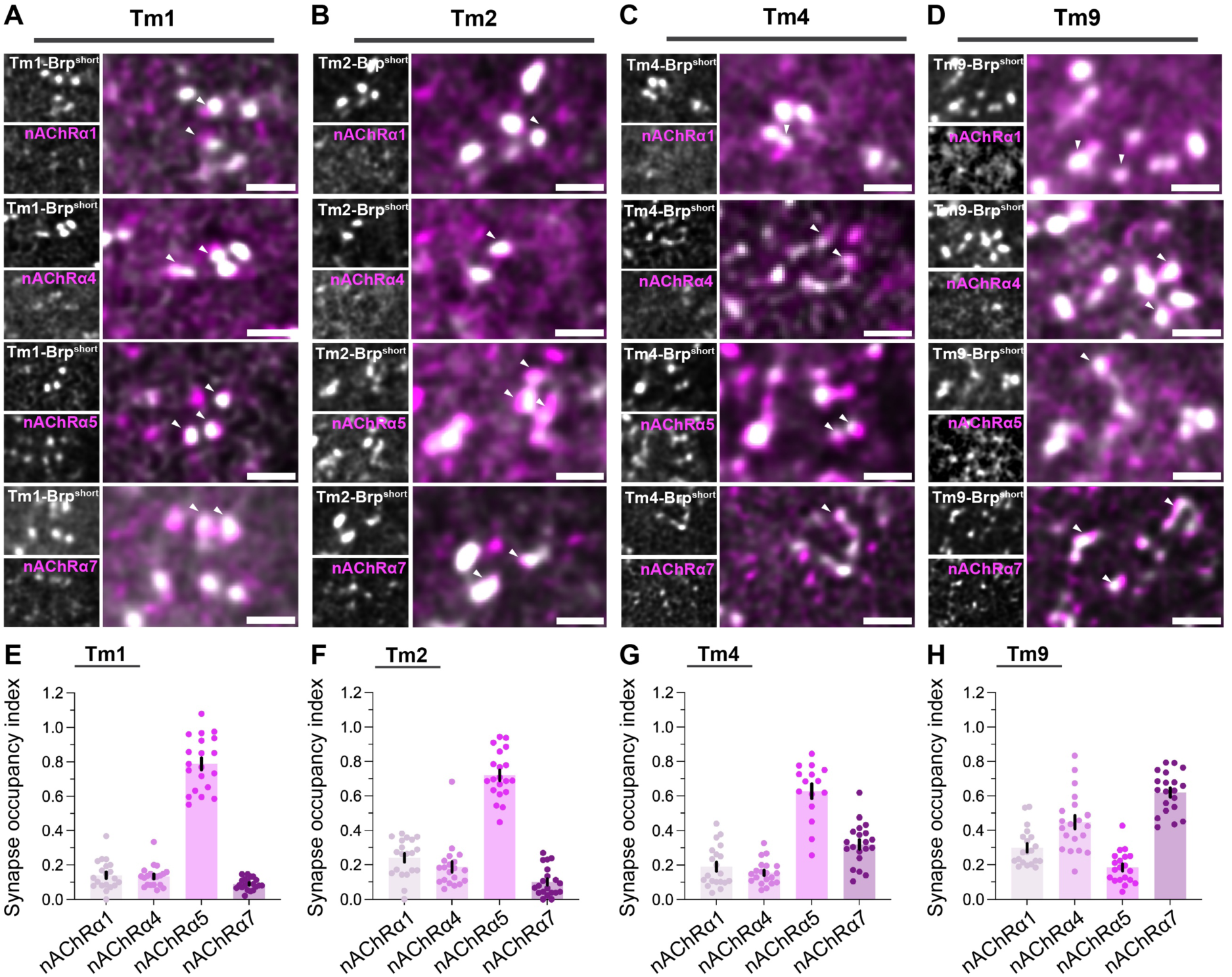
Differential synaptic localization of nAChRα subunits in Tm synapses. (A-D) Trans-synaptic alignment of Tm1, Tm2, Tm4 and Tm9 boutons (Tm-Gal4> UAS-Brp^short::mStraw^, grey) with the nAChRα1, nAChRα4, nAChRα5 and nAChRα7 subunits (nAChRα::EGFP, magenta) in LO1. Arrows indicate the Brp^short::mStraw^-receptor punctum trans-synaptic alignment. Scale bar 1μm. (E-H) Synapse occupancy of Tm1, Tm2, Tm4 and Tm9 boutons with nAChRα1 (pale magenta), nAChRα4 (light magenta), nAChRα5 (magenta) and nAChRα7 (dark magenta) subunits in LO1 (optic lobes n=4, ROIs N=20, for Tm4-nAChRα5 n=3, N=15). Data is mean ± SEM.

Whether Tm1, Tm2, Tm4 and Tm9 cells differ from one another by differentially using muscarinic receptors as well, is currently unknown. Overexpression of the mAChR-B type in T5 cells led to the mAChR-B localization in Tm9-T5 synapses (Figure S9A), which was more evident when compared to the nAChRα5 subunit (Figure S9B). The apparent lack of a generalized cellular mistargeting of the mAChR-B overexpression line (Figure S2I), does not exclude the existence of synaptic mistargeting events. Consequently, the Tm9-mAChR-B results are indicative rather than definitive. Overall, our nAChRα subunit mapping across Tm synapses in LO1 revealed that Tm1, Tm2 and Tm4 cells differ from Tm9 at the postsynaptic level^48,57^.

### Spatial organization of Tm-nAChR**α** synapses in T5 dendrites

The compartmentalization of inputs and subsequently of receptors on T5 dendrites is essential for the T5 direction-selectivity^53,54^. Thus far, receptor localization in Tm-to-T5 synapses has been addressed by combining single-cell receptor localization and neuronal input spatial wiring information^60,79^. To advance this, we analyzed both T5a and T5c dendrites as representatives of the two directional systems from the FAFB^62^ dataset via the FlyWire (flywire.ai)^63,64^ interface. Manual proofreading of predicted synapses^80,81^ revealed the highest number of Tm9 synapses, compared to Tm1, Tm2 and Tm4, in both T5 subtypes as previously reported^40,51^ (Figure 4A,B). To account for these Tm-to-T5 differences in synapse counts, we measured the synapse occupancy index (Figure 3E-H). Next, we sought to identify the spatial wiring of the four Tm neurons on T5 dendrites. Tm1, Tm2, Tm4 and Tm9 synapses localize to the central dendritic compartment, with Tm9 synapses extending to the distal compartment as well, as previously shown^40,51^ (Figure 4C,D). Together with our subunit synaptic localization results, we conclude that the nAChRα1, nAChRα4 and nAChRα7 subunits localize across the dendrite to Tm1, Tm2, Tm4 and Tm9-T5 synapses, while the nAChRα5 subunit localizes primarily in Tm1, Tm2 and Tm4-T5 synapses in the central compartment of T5 dendrites (Figure 4E).

**Figure 4.**
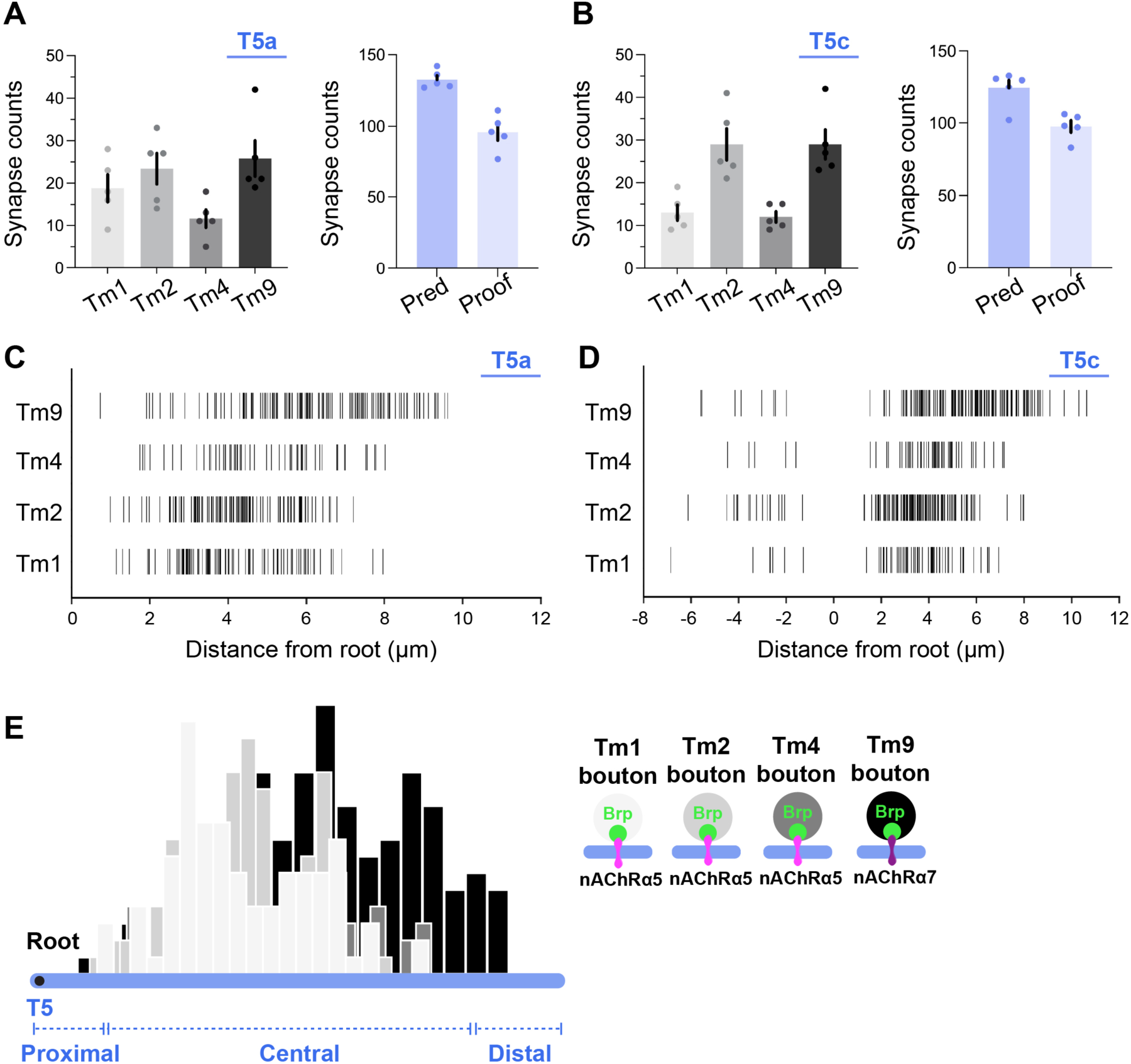
Wiring of Tm1, Tm2, Tm4 and Tm9 on T5 dendrites. (A) Counts of Tm1, Tm2, Tm4 and Tm9 synapses in T5a dendrites (T5a cells n=5, flywire.ai) (left). Counts of predicted (n=663) to proofread (n=479) Tm synapses in T5a dendrites (right). Data is mean ± SEM. (B) As in (A) for T5c dendrites (T5c cells n=5, flywire.ai) (left). Counts of predicted (n=621) to proofread (n=488) Tm synapses in T5c dendrites (right). Data is mean ± SEM. (C) Dendritic localization of Tm synapses across the T5a anterior-posterior axis (T5a cells n=5). Root is considered the first dendritic branch point. (D) Dendritic localization of Tm synapses across the T5c dorso-ventral axis (T5c cells n=5). (E) Schematic representation of Tm-T5a dendritic synapses and the observed postsynaptic receptor heterogeneity in terms of the most abundant nAChR subunit as in Figure 3E-H.

### AChRs differentially affect the optomotor response

When stimulated by rotatory visual motion, flies perform corrective steering maneuvers in their flight or walking trajectories. This behavior is called the optomotor response^30,82,83^ and T4 and T5 cells are indispensable for its manifestation in the ON and the OFF pathway accordingly^84–87^. Therefore, molecular alterations that are crucial for the proper T5 function should be reflected in the fly’s optomotor response behavior. We used reversed genetics to test the role of AChRs in the OFF-motion pathway. Using RNAi knock-down, we tested whether nAChRα1, nAChRα4, nAChRα5, nAChRα7 and mAChR-B were required for the optomotor response on the ‘fly-on-ball’ set up. Given that each receptor has a unique developmental onset of expression (Figure S1D), we used a T4-T5 driver line that was active from the third larval stage onwards^84,88^ and we raised flies in 29°C^89,90^ in order to achieve a strong knock-down of protein levels. The driver line targeted all T5 subtypes and allowed for the RNAi induction prior to the peak of AChR expression (Figure S3G,H). Finally, we confirmed the RNAi-induced reduction in AChR subunit expression with immunohistochemistry (Figure S3A-F).

Flies walking on an air-suspended ball were exposed to a set of OFF edges moving in sixteen directions, which elicit a typical profile of angular walking velocities, with the highest amplitudes to horizontal motion and lowest amplitudes to vertical motion^45^ (Figure 5A). Knock-down of the nAChRα1 subunit did not alter the angular velocity response profile to the OFF edges (Figure 5B,G). Knock-down of the nAChRα4 subunit or the mAChR-B type resulted in an increase of the angular velocity in response to OFF edges moving in both horizontal directions (Figure 5C,F,G). In contrast, knock-down of the nAChRα5 and the nAChRα7 subunits led to a decrease in the angular velocity amplitude in response to horizontal moving edges (Figure 5D,E,G). The nAChRα3 and nAChRβ1 subunits were recently found to localize on T5 dendrites^60^, while the mAChR-A type exhibited high RNA expression levels^43,58^ and expression in T4-T5 cells based on our enhancer-trap screen (Figure S1C,D, S4K). Knock-down of nAChRα3, nAChRβ1 and mAChR-A did not affect the angular velocity profile in response to OFF edges (Figure S10A-E). However, the RNAi efficiency against the nAChRα3, nAChRβ1 and the mAChR-A could not be verified. Therefore, these AChRs should not be excluded from playing an important role in the optomotor response. In summary, we found that AChRs play different roles in the optomotor response.

**Figure 5.**
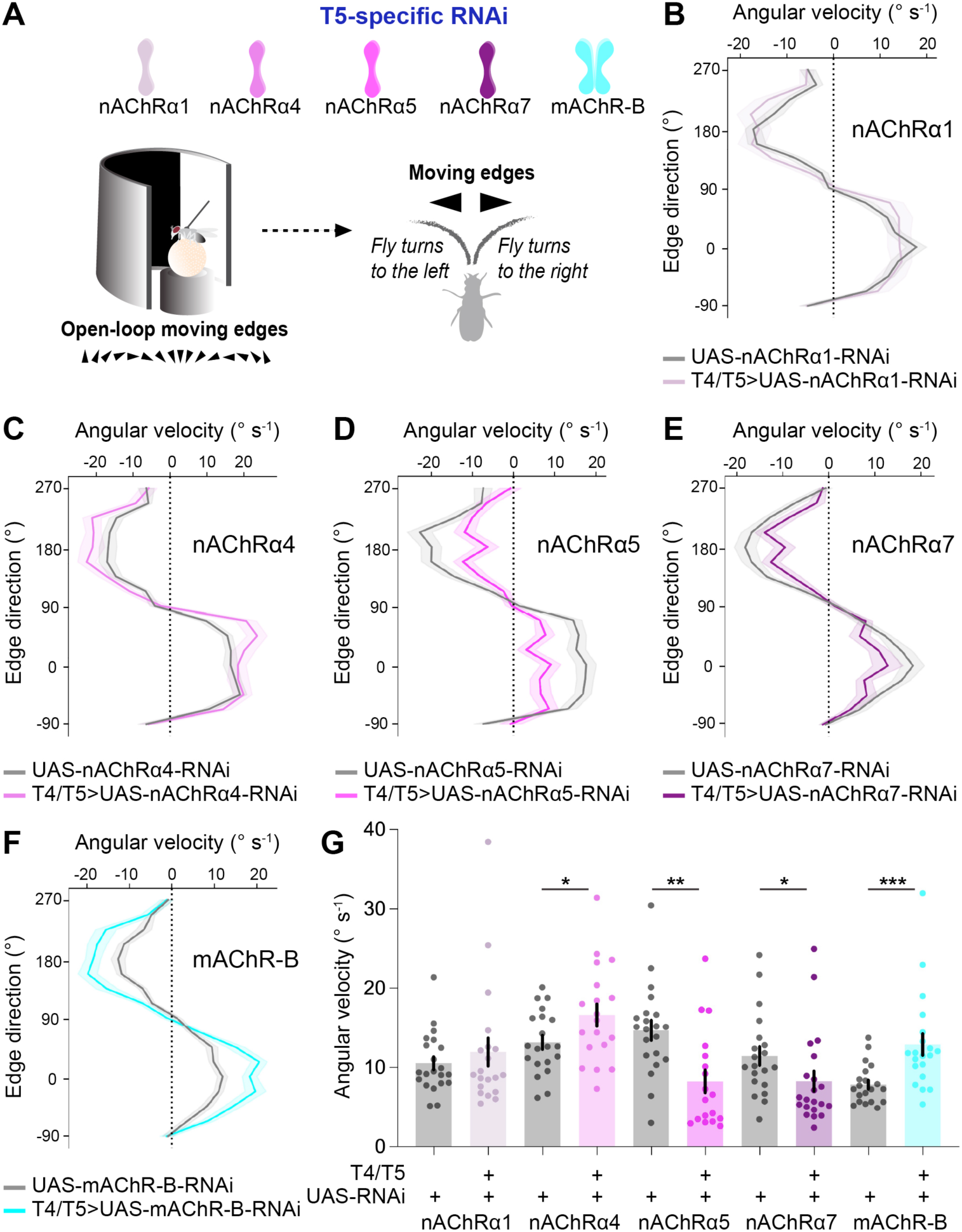
Fly optomotor response post-AChR knock-down. (A) Schematic workflow representation for the behavioral assessment of AChR knock-down flies. (B) Trajectory (angular velocity) of nAChRα1 knock-down flies (R42F06-p65.AD; VT043070-Gal4.DBD>UAS-nAChRα1-RNAi, pale magenta, flies n=20) while presented to a sequence of upward to rightward to downward to leftward OFF edge motion compared to the parental control flies (UAS-nAChRα1-RNAi, grey n=21). The dark lines represent the mean ± SEM. (C-F) Same as in (B) for the nAChRα4 (light magenta n=19, grey n=20), nAChRα5 (magenta n=18, grey n=21), nAChRα7 (dark magenta, n=21, grey n=20), mAChR-B (cyan n=20, grey n=20) knock-down flies. (G) Angular velocity of control and AChR knock-down flies. The normality of distribution was assessed with the use of Shapiro-Wilk test. *p<0.05, **p<0.01, ***p<0.001. Data is mean ± SEM.

### Directional tuning of T5 cells changes after AChR knock-down

The nAChRα5 and nAChRα7 subunits localize both on T5 dendrites and in T5 axonal terminals (Figure S2K). The effects on the behavioral responses to OFF-edge horizontal motion that were observed upon nAChRα5 and nAChRα7 knock-down (Figure 5D,E,G) could either be due to functional roles in T5 dendrites or in T5 axonal terminals. To discern between these two possibilities, we measured calcium responses in T5 axonal terminals. We performed two-photon calcium imaging in T5c neurons, which are sensitive to upward motion as the preferred motion direction^34^. nAChRα4, nAChRα5, nAChRα7 or mAChR-B knock-down flies were stimulated by dark edges moving in sixteen different directions, while calcium responses were monitored in T5c axonal terminals in layer 3 of the lobula plate (LOP3) (Figure 6A, Figure S11A-C). As in the behavioral experiments, RNA interference was induced prior to peak receptor expression with the use of a T4-T5 promoter that is active from the third larval stage^88^, and raising flies in 29°C^89,90^. Knock-down of the nAChRα4 subunit led to a slight increase in calcium responses in response to the non-preferred direction (270°) of T5c neurons (Figure 6B). Similarly, knock-down of the nAChRα5 subunit resulted in higher calcium responses to the edge moving in the non-preferred direction (Figure 6C), suggesting nAChRα5, and to a lesser extent, nAChRα4 contribute to directional tuning in T5 neurons. Knock-down of the nAChRα7 subunit resulted in an increase in calcium responses to both the null and preferred motion direction (Figure 6D). Knock-down of the mAChR-B caused an overall increase of calcium T5 responses across all presented stimuli directions (Figure 6E). Individual fly responses shown in Figure S11D, show that the nAChRα4 and nAChRα5 subunit knock-down flies show high inter-fly variation, whereas the nAChRα7, mAChR-B and mCherry RNAi flies exhibit more stable response patterns. In general, the nAChRα4, nAChRα5 subunits and mAChR-B type significantly influenced the T5 directional tuning (Figure 6F). Together these experiments show that an interplay of nicotinic and muscarinic receptors shapes the T5 directional responses.

**Figure 6.**
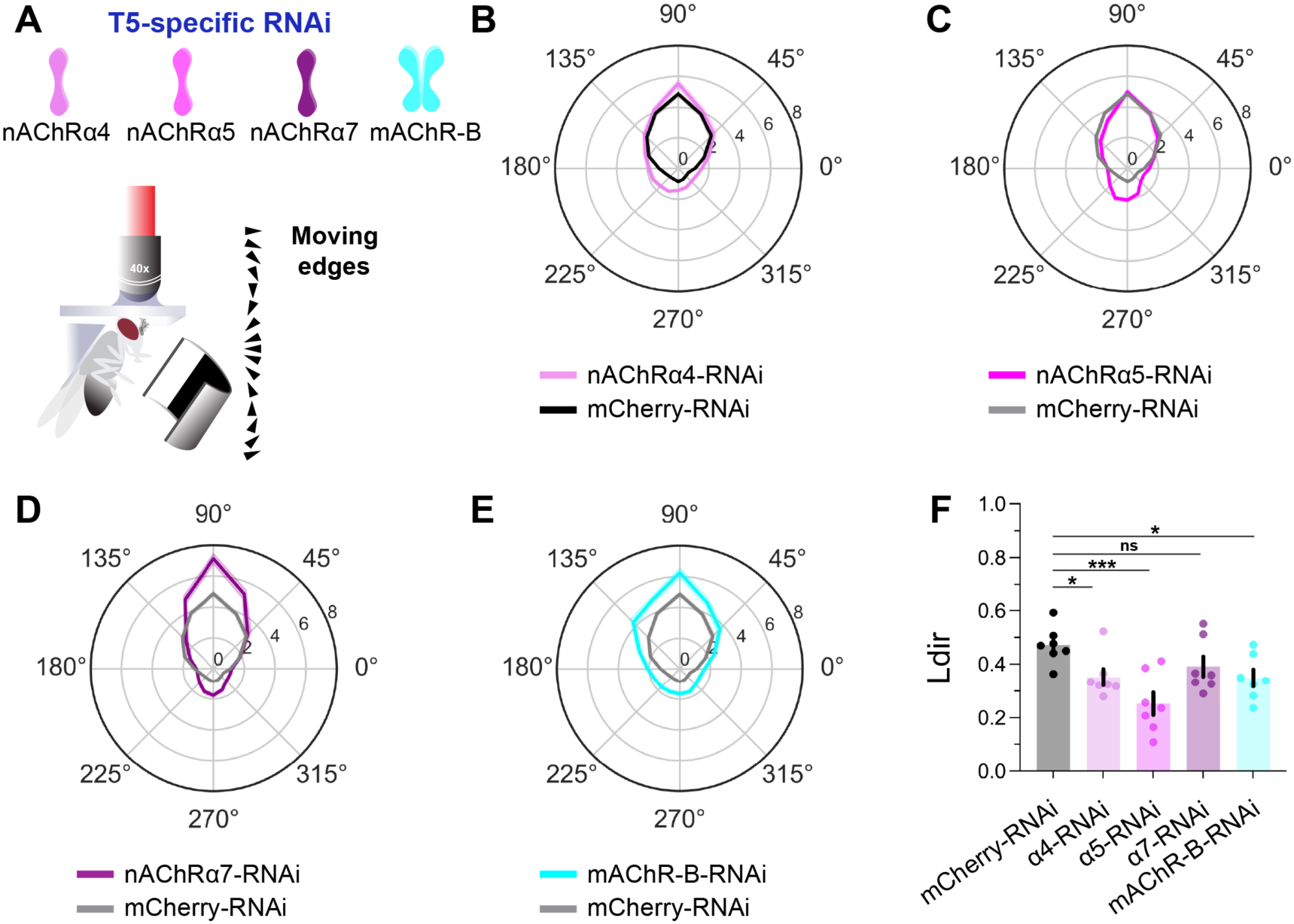
Directional tuning of T5 cells post-AChR knock-down. (A) Schematic workflow representation for the two-photon calcium imaging in AChR knock-down T5 cells. (B) Polar plot of the maximum responses (ΔF/F) across sixteen OFF edge directions (°) of T5c cells for mCherry controls (black, flies n=7, ROIs N=85, shown as grey in C, D and E) and nAChRα4 RNAi (light magenta, n=7, N=61) conditions (R39H12-Gal4>AChR-RNAi, VT50384-LexA>LexAop-IVS-GCaMP6m). The dark lines represent the mean (68% CI). (C) Same as in (B) for nAChRα5 RNAi flies (magenta, n=7, N=82). (D) Same as in (B) for nAChRα7 RNAi flies (dark magenta, n=7, N=60). (E) Same as in (B) for mAChR-B RNAi flies (cyan, n=7, N=54). (F) Directional tuning index (Ldir). Flies n=7. The normality of distribution was assessed with the use of Shapiro-Wilk test. *p<0.05, ***p<0.001. Data is mean ± SEM.

Cholinergic neurotransmission in T5 cells takes place in two distinct locations: on the dendrite in LO1, where T5-T5 dendro-dendritic synapses^40^ reside, and on the axonal terminal niche in LOP3, where T4-T5 and T5-T5 axo-axonic synapses reside^91^. Calcium responses in nAChRα5 subunit knock-down flies exhibited different profiles in LO1 and LOP3 (Figure S12A-C). Calcium responses in T5 dendrites were of lower amplitude, while responses from T5 axonal terminals increased in response to the non-preferred direction (270°) in nAChRα5 knock-down compared to control mCherry RNAi flies (Figure S12B,C). Such differences in calcium responses imply distinct roles of AChRs in the two cholinergic neurotransmission niches of T5 neurons.

## Discussion

In this study we investigated the cholinergic neurotransmission onto T5 neurons of *Drosophila* and found that T5 cells express a variety of AChRs on their dendrites, which are surprisingly differentially distributed across the cholinergic Tm1-, Tm2-, Tm4-, Tm9-to-T5 synapses. This acetylcholine receptor variety translated into a functional variety, as portrayed by both the fly optomotor response and the T5 directional tuning.

In mammals, nicotinic AChRs comprise nine α subunits (α1-7,9,10), four β subunits (β1-4), one γ, one δ and one ε subunits^92^, whilst muscarinic AChRs consist of five M (M1-5) types^93^. This AChR variety is also found, but to a lesser extent, in *Drosophila melanogaster*, with its ten nAChR subunits and three mAChR types. Does this structural variety allow for differences in neuronal function? The nAChRα5 subunit was found to be required in M4/6 mushroom body output neurons (MBONs) for coding immediate appetitive memories^22^. The mAChR-A type was observed on the dendrites of Kenyon cells, mediating aversive olfactory learning in adult flies^14^. Moreover, a study of the simple visual neural circuit in *Drosophila* larvae showed that the glutamatergic interneuron glu-lOLP mediates OFF detection via mAChR-B signaling^23^, while the interplay of nAChRs and mAChRs in regulating acetylcholine modulation of larval locomotion has been pharmacologically assessed with bath application of AChR agonists and antagonists^15^. Our study provides another example of diverse AChR functions in neuronal circuits, where an interplay between nicotinic and muscarinic receptors is observed in the visual system of the adult fly.

### AChR expression on T5 dendrites

For a long time, the expression of AChRs in the visual system of the fruit fly has been under investigation^60,79,94–96^. Advancements in genomics and molecular engineering now allow access to the AChRs of interest by endogenously tagging them with fluorescent proteins^22,60,79^. This allows for qualitatively and quantitatively assessing the AChR neuronal expression, and avoiding the ambiguity that was introduced via combinatorial studies of electrophysiology and antagonist application in terms of qualitative AChR identification^97^. We identified the nAChRα1, nAChRα4, nAChRα5 and nAChRα7 subunit protein localization on T5 dendrites (Figure 1). The nAChRβ1 subunit has recently been found to co-localize with the nAChRα1 subunit on T5 dendrites^60^. Whether this co-localization depicts a pentameric co-organization as previously speculated in lobula^94^, remains to be proven, but it certainly indicates the stoichiometrical nAChR complexity^98,99^ that exists in the optic lobe. For the mAChR-mediated cholinergic neurotransmission, our results together with previous RNA-seq studies^43,58,59^ indicate the expression of the mAChR-B type in T5 cells.

### nAChR**α** subunits in Tm-to-T5 synapses

Sparse neuronal labelling allowed us to verify the explicit genetic access to Tm1, Tm2, Tm4 and Tm9 neurons, which then could be combined with synaptic labelling by Brp. While mapping the most highly expressed subunits nAChRα1, nAChRα4, nAChRα5 and nAChRα7 across Tm1, Tm2, Tm4 and Tm9 synapses in LO1, we found the nAChRα5 subunit preferentially localizing to Tm1, Tm2, Tm4 rather than Tm9 synapses, implying that the functional differences observed among Tm1, Tm2, Tm4 and Tm9 cells^48,57^ are extended to the postsynaptic receptor level. Tm9 is functionally different from Tm1, Tm2 and Tm4 by acting as a low-pass filter^48,49,57^ and Tm9-to-T5 synapses use a different nAChR set for their cholinergic neurotransmission. This suggests that the preferred direction enhancement in T5 cells might not only be shaped by the different time-courses of the input signals they receive, but it additionally relies on postsynaptic molecular mechanisms. Moreover, the mAChR-B localization in Tm9-T5 synapses, together with the nAChRα1, nAChRα4, nAChRα7 and to a lesser extend nAChRα5 localization, points towards an interplay between ionotropic and metabotropic receptors. The role of such receptor interactions is still unclear, but recent examples indicate that it might potentiate the nicotinic AChR responses^20^.

Attempts to test the role of neuronal input compartmentalization onto T5 dendrites has led to combinatorial approaches, where the spatial wiring of inputs together with the receptor dendritic localization were addressed. Together with previous reports^40,51^, our EM analysis shows a spatial overlap of Tm synapses on T5 dendrites, specifically on the central dendritic compartment. It is therefore risky to draw conclusions about receptor localization in Tm-to-T5 synapses via single-cell receptor localization experiments, especially due to the shared cholinergic nature of the input neurons and their spatial overlap. In this study, we introduce a new methodology, where EM-derived neuronal input spatial wiring and synaptic receptor mapping are combined to extract pre-, post-synaptic and spatial wiring information. Ultimately, single synapse-to-single dendrite approaches will improve the resolution of connectivity studies, leading to a T5 AChR-connectome^100^.

### AChRs roles in the OFF-motion pathway

Do these individual nAChR subunits and mAChR types differentially control T5 motion detection? We found that both ionotropic and metabotropic AChR signaling contribute to T5 function at the behavioral and neuronal activity levels, implying that an unforeseen complexity of cholinergic neurotransmission likely underlies motion detection in the OFF pathway. Notably, the nAChRα5 and nAChRα7 localization in T5 axonal terminals in addition to dendrites, might result in behavioral alterations deriving from other than dendritic computations. As the behavioral output involves many T5 downstream partners^83,101,102^, compensatory mechanisms might also be in place, potentially masking immediate effects in T5 cells.

Calcium imaging in T5c neurons revealed an increase in the null direction responses among all nAChRα4, nAChRα5, nAChRα7 and mAChR-B knock-down flies. In contrast, the nAChRα7 or mAChR-B knock-down led to amplified preferred direction responses. We expected that the nAChRα5 subunit knock-down, as being the most highly expressed subunit along Tm1-, Tm2-, Tm4- and Tm9-T5 synapses, would have severely attenuated the T5 preferred direction responses. However, this was not the case, suggesting an important role for the nAChRα1, nAChRα4 and nAChRα7 subunits in preserving Tm1-, Tm2-, Tm4- and Tm9-to-T5 function. Lastly, the mAChR-B knock-down-induced effects resemble that of a raised inhibition, which align with the receptor’s reported inhibitory role^23^. Our results show that there is an interplay of nicotinic and muscarinic AChRs taking place on T5 dendrites. The distinct stoichiometries as well as the ionic properties of the different nAChR subunits need to be identified in the future to help us better understand the molecular mechanisms underlying direction selectivity in T5 cells.

### Limitations of this study

Multiple aspects should be taken into account when interpreting RNA interference experiments. Firstly, an AChR subunit compensation^103^ might take place, concealing the true receptor subunit function. To which extent such a mechanism is activated in T5 neurons, and towards which subunits, is currently unknown. Secondly, the imaging location (T5 dendrites in LO1 versus T5 axonal terminals in LOP) should be considered. The T5-T5 dendro-dendritic synapses, the T5-T5 and T4-T5 axo-axonic synapses^91^ are cholinergic. Our RNA interference of nAChRα5 and nAChRα7 subunits will affect such synapses as well, which might confound our conclusions on T5 dendrites. In an effort to attend to this, we observed different responses in T5 dendrites compared to T5 axonal terminals. The difference in dendrite and axon responses could result from differences in the density of ionotropic AChRs and other voltage gated calcium channels between the two T5 compartments or from an uncharacterized influence of T5-T5 dendro-dendritic and T5-T5 and T4-T5 axo-axonic synapses (Figure S12). Finally, even though we used standardized conditions for our RNAi induction experiments, fly-to-fly variability was encountered in nAChRα4 and nAChRα5 knock-down flies (Figure S11). This led us to believe that the RNAi efficiency may differ across individuals and compensation mechanisms at the individual level might be activated.

## ACKNOWLEDGMENTS

The authors thank D. Owald, C.H. Lee, F. Pinto-Teixeira, the Bloomington Drosophila Stock Center, the Vienna Drosophila Stock Center and the Developmental Studies Hybridoma Bank for flies and antibodies. We are grateful to the Princeton FlyWire team and members of the Murthy and Seung labs for development and maintenance of FlyWire (supported by BRAIN Initiative grant MH117815 to Murthy and Seung). We are thankful to B. Zuidinga for technical assistance with behavioral experiments and calcium imaging analysis; M. Sauter for technical assistance with fly dissections; S. Prech for technical assistance with experimental setups; R.M. Vieira and J. Pujol-Martí for discussions. E. Samara is a member of the Graduate school of Systemic Neurosciences (GSN) Munich. I.M.A.R. was funded by LMU and the DFG grant nr RI 3644/2-1. This work was funded by the Max Planck Society.

## AUTHOR CONTRIBUTIONS

E.S. and A.B. conceived this study. E.S., T.S. and I.M.A.R. designed the experiments. E.S. performed all experiments with the help of T.S. and M.B.L. T.S. contributed to the receptor synaptic localization experiments and analysis. M.B.L contributed to the optomotor response experiments. J.H. contributed to the calcium imaging analysis. E.S. wrote the manuscript with the help of I.M.A.R. and A.B. All authors have read and commented on this manuscript.

## DECLARATION OF INTERESTS

The authors declare no competing interests.

## STAR⍰Methods

## RESOURCE AVAILABILITY

### Lead contact

Further information and requests for data should be directed to and will be fulfilled by the Lead Contact, Alexander Borst.

### Materials availability

The study generated a new fly line and is available from lead contact upon request.

### Data and code availability

- This paper analyzes existing, publicly available data from FlyWire (flywire.ai).
- Data and code for analysis are publicly available at the at the Edmond Open Research Data Repository of the Max Planck Society.
- Any additional information required to reanalyze the data reported in this paper is available from the lead contact upon request.

## EXPERIMENTAL MODEL AND SUBJECT DETAILS

### Fly husbandry

All flies were raised on corneal agar medium under standard conditions (25°C, 60% humidity, 12h light/dark cycle). For the interference experiments (RNAi), flies (control and RNAi) at early pupa stages were transferred from 25 to 29°C. For the Multi-color FlpOut experiments, flies at early pupa stages were heat-shocked for 30 minutes at 37°C.

## METHOD DETAILS

### Generation of UAS-CRASP line

The pQUAST-syb-spCFP1-10 vector^72^ and the pJFRC7-20XUAS-IVS-mCD8::GFP vector^104^ were digested with XhoI and XbaI restriction enzymes and the syb-spCFP1-10 DNA fragment was extracted and T4-ligated in the pJFRC7 vector. The AGGCACACCGAAACGACTAA and CAGTTCCATAGGTTGGAATCTAAAA primers were used for sequencing the successful insertion of the DNA fragment. Embryo injections were performed by BestGene Inc (Chino Hills, CA, USA) in the VK00027 attp landing site.

### RNAi experiments

To ensure high interference efficiency, given that both long dsRNA (nAChRα1, nAChRα4 and nAChRβ1) and short shRNA (nAChRα3, nAChRα5 and nAChRα7, mAChR-A, mAChR-B) were used, flies were raised (at early pupa stages) at 29°C^89,90^ and the RNAi was induced prior to the receptor expression peak. Dicer was not used for the long dsRNA experiments as previous studies^22,105,106^ showed positive AChR-RNAi induction without dicer co-expression. Our immunohistochemical screening spoke in favor of the nAChRα1, nAChRα4, nAChRα5, nAChRα7 and mAChR-B RNAi effectiveness (Figure S3).

### Immunohistochemistry

Fly brains (aged 2-5 days, for G-and C-RASP experiments aged 10-12 days) were dissected in cold phosphate-buffered saline (PBS) and fixed in 4% paraformaldehyde (in PBS with 0.1% Triton X-100) for 24 minutes at room temperature, followed by three 10-minute washes in PBT (PBS and 0.3% Triton X-100). Brains were then incubated with primary antibodies in PBT for 48 hours at 4°C. After being incubated for 2 hours at room temperature, brains were washed four times for 15 minutes each in PBT and then incubated with secondary antibodies diluted in PBT for 48 hours at 4°C. After being incubated for 2 hours at room temperature and four 15 minutes PBT washes, brains were mounted for immediate sample viewing. For the subunit mapping (Figure 3), brains were blocked in 5% normal goat serum (NGS) in PBT for 1 hour at room temperature and then incubated with the primary and secondary antibodies for 24 hours instead of 48 hours.

### Confocal microscopy

Images were acquired in a Leica Stellaris 5 laser scanning confocal microscope with a 20x glycerol 0.75 NA HC Plan-apochromat and 63x glycerol immersive 1.3 NA HC Plan-apochromat objective at 2048 x 2048 x 0.4μm image resolution. The deconvolution setting (Lightning) of this microscope was used for the acquisitions in Figure 3. Image analysis was performed with Fiji (ImageJ)^107^. For the acquisitions in Figure S4, a Leica SP8 laser scanning confocal microscope with a 63x oil immersive 1.3 NA HCX Plan-apochromat objective was used at 2048 x 2048 x 0.4μm image resolution.

### Behavior

To avoid behavioral effects introduced by sexual dimorphism^26^, only female flies, 1-3 days old, were used in our experiments. Flies were attached to a pin post-cold anaesthetization with the use of light-curing glue and dental curing light (440 nm, New Woodpecker). Each locomotor recorder, out of the four that were simultaneously used, consisted of an air-suspended sphere (6mm in diameter and 40 mg weight) made of polyurethane foam and coated with polyurethane spray^85,108^. A constant airflow by a rotary vane pump (G6/01-K-EB9L Gardner Denver Thomas) allowed for the free rotation of the sphere on the ball-shaped sphere holder. The sphere’s rotation was tracked by two optical tracking sensors, focusing two 1-mm^2^ equatorial spots at ±30° from the center of the infrared LED-illuminated (JET-800-10, Roithner Electronics) sphere. Rotational data were tracked at 4 kHz and digitized at 200 Hz^109^. To achieve a successful fly positioning on the sphere, a camera (GRAS-20S4M-C, Point Grey Research) was used. A custom-made Peltier system (QC-127-1.4-6.0MS, Quick-Cool) controlled the airflow temperature at 34 ± 0.1°C, based on the readouts deriving from a thermometer placed just below the sphere, so as to ensure prolonged walking. A U-shaped visual arena comprising of three 120-Hz LCD screens (Samsung 2233 RZ) (reaching a maximum luminance of 131 cd m^−2^), allowed for the presentation of visual stimuli spanning approximately 270° in azimuth and 120° in elevation of the fly’s visual field at a resolution of <0.1°. Panda3D was used in Python v.2.7 for the visual stimulus generation^85^.

### Two-photon microscopy

For functional imaging experiments, we used a custom-build two photon laser scanning microscope equipped with a 40x water 0.80 NA IR-Achroplan objective (Zeiss)^34^. Flies (1-3 days) were cold-anaesthetized and the legs and thorax were glued on a Plexiglas holder with the use of light-curing glue. The head was bend down, fitted in an aluminum opening and the holder was clamped in a recording chamber. Saline solution (PBS; 137 mM NaCl, 3 mM KCl, 8 mM Na_2_HPO_4_, 1.5 mM KH_2_PO_4_, pH 7.3) was introduced and the posterior side of the fly’s optic lobe was exposed by a small incision of the head. Muscles, adipose tissue and trachea were manually removed. Images were acquired at a 64 x 64-pixel resolution and a frame rate of 15□Hz in Matlab R2013b (MathWorks) using ScanImage 3.8 software (Vidrio Technologies, LLC).

### Visual stimulation

For the behavioral experiments, OFF edges (50% contrast) moved at 16 evenly spaced directions with a velocity of 60° s^−1^, were randomized and crossed the whole arena span within 5s. Each experiment lasted for ∼55 min, including 35 trials of OFF edges, of which the first 15 trials were excluded from analysis, as they were appointed to temperature stabilization and fly accommodation. The same inclusion criteria as in a previous study^45^ were used.

For the calcium imaging experiments, we used a custom-built projector-based arena^57^, where two micro-projectors (TI DLP Lightcrafter 3000) projected visual stimuli, with a refresh rate of 180 Hz and maximum luminance of 276 ± 48 cd/m^2^, onto the back of a cylindrical screen. The arena covered 180° in azimuth and 105° in elevation of the fly’s visual field. OFF edges (92% contrast) moved at 16 evenly spaced directions with a velocity of 30° s^−1^, were randomized and repeated four times.

### Trans-synaptic alignment

To assess the trans-synaptic alignment between the receptors and the synapses of interest, we calculated the fluorescence intensity values (later normalized to the maximum pixel intensity value) in a 1 μm line positioned across the synapse punctum in a single z plane using Fiji-ImageJ^107^. Data analysis was performed with Python v.3.9.18 with the use of seaborn 0.12.2, pandas 1.5.3 and numpy 1.23.3.

### EM analysis

For the Tm-T5 wiring, we used the FAFB volume^62^ via the FlyWire (flywire.ai) proofreading environment^63,64^ and chose five reconstructed T5a and five T5c neurons from the right optic lobe (Table S3). T5 and Tm1, Tm2, Tm4, Tm9 identities were verified by users’ annotations and by comparison with previous electron microscopy^40^ and our light microscopy reconstructions. Synapse number Buhmann predictions^80,81^ between the T5 neuron of interest and its respective input neurons were acquired from FlyWire’s connectivity viewer. Cleft score was set to 50, so as to eliminate synaptic redundancy resulting from the combination of synaptic connections between the same neurons when their presynaptic locations are within 100nm^2^. For each synapse, we sought four morphological markers: a. synaptic vesicles, b. the protein dense T-bar structure, c. synaptic cleft, and d. postsynaptic densities (or postsynaptic domains). Every synaptic locus was assessed in a volume of five up to seven brain slices, 40nm each. Only synapses that displayed all four morphological markers were included in our dataset. We treated proofread synapses as points in T5 dendrite space by collecting the x,y,z coordinates of each T-bar (pre-synapse). Synapse numbers in this study correspond to manually proofread synapses.

For synapse distributions across the anterior-posterior and dorso-ventral dendritic arborization axis (Figure 4C,D), we measured the Euclidean distance of every synapse from the root (first branching point) of the dendrite. Distances in μm were calculated by multiplying the y and z coordinates of each T-bar with 4 and 40 accordingly (voxel=4×4×40nm). To correct for anterior-posterior and dorsal-ventral dendrite length differences, we normalized each synapse-to-root Euclidean distance to the longest T5a and T5c dendrite.

For the identification of Tm1, Tm2, Tm4 and Tm9 synaptic partners in LO1 (Figure S8), we used Codex^62,63,64^. We searched for all the Tm output neurons in five Tm1 (FlyWire ID: 720575940620976493,720575940608883465,720575940623583428, 720575940632600647,720575940621146733), Tm2 (FlyWire ID: 720575940632142904,720575940640453437,720575940640230259,720575940630551670,720575940622364961), Tm4 (FlyWire ID: 720575940637976666, 720575940620745163, 720575940621582401, 720575940626612228, 720575940620782171) and Tm9 (FlyWire ID: 720575940613374873, 720575940620814356, 720575940609254851, 720575940626242348, 720575940623771061) cells, and selected only those outputs whose Tm synapses were restricted in LO1 (excluding all Tm medullary output neurons). For the output neurons that only formed connections with Tm neurons in LO1, we used the predicted synapse number, but for those output neurons that made Tm connections in other LO layers as well, we manually proofread the LO1 synapses. The reconstructed neurons for Figure 2 correspond to FlyWire IDs 720575940621338038, 720575940614527742, 720575940616041926 and 720575940616813142 for Tm1, Tm2, Tm4 and Tm9 respectively.

## QUANTIFICATION AND STATISTICAL ANALYSIS

### nAChRα density

Receptor puncta that localized in T5 dendrites were quantified along 8μm x 8μm x 4μm voxels in Fiji-ImageJ^107^. The total number of puncta divided to the analyzed LO1 volume resulted in the nAChRα density.

### Synapse occupancy

The total number of Brp-receptor co-localizing puncta in voxels of 8μm x 8μm x 4μm in Fiji-ImageJ^107^, was divided to the number of Brp puncta, resulting in the synapse occupancy index.

### RNAi efficiency

The mean grey value of ten 8μm x 8μm ROIs per optic lobe in LO1 and LOP (layer 1 and 2) was measured via Fiji-ImageJ^107^ and was normalized by subtracting the mean grey value of one 8μm x 8μm ROI in the inner optic chiasm.

### Behavior

The optomotor response was quantified as the absolute average rotational velocity during 5 s of edge motion in each direction. Data analysis was performed with Python v.3.9.18 using seaborn 0.12.2, pandas 1.5.3 and numpy 1.23.3.

### Calcium imaging

Calcium imaging data were analyzed as described in^57^ with a custom written software in Python v.2.7. ROIs were drawn manually across LO layer 1 and LOP layer 3. Only ROIs that repeatedly responded to the given visual stimulus and exhibited consistent-across the four repetitions-responses, were included in the dataset. For each ROI, the time courses of relative fluorescence changes (ΔF/F) were calculated and responses to stimulus were baseline-subtracted and averaged over repetitions. Maximum responses were aligned to 90°. The directional tuning index (Figure 6F, Ldir) was calculated as the magnitude of the resultant vector divided by the sum of the individual vectors’ magnitudes:

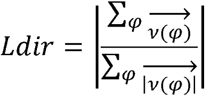

### Statistical analysis

Detailed analysis is reported in figure legends and table S2 and was performed in GraphPad Prism v.9.3.0.

## SUPPLEMENTAL INFORMATION TITLES AND LEGENDS

**Figure S1.**
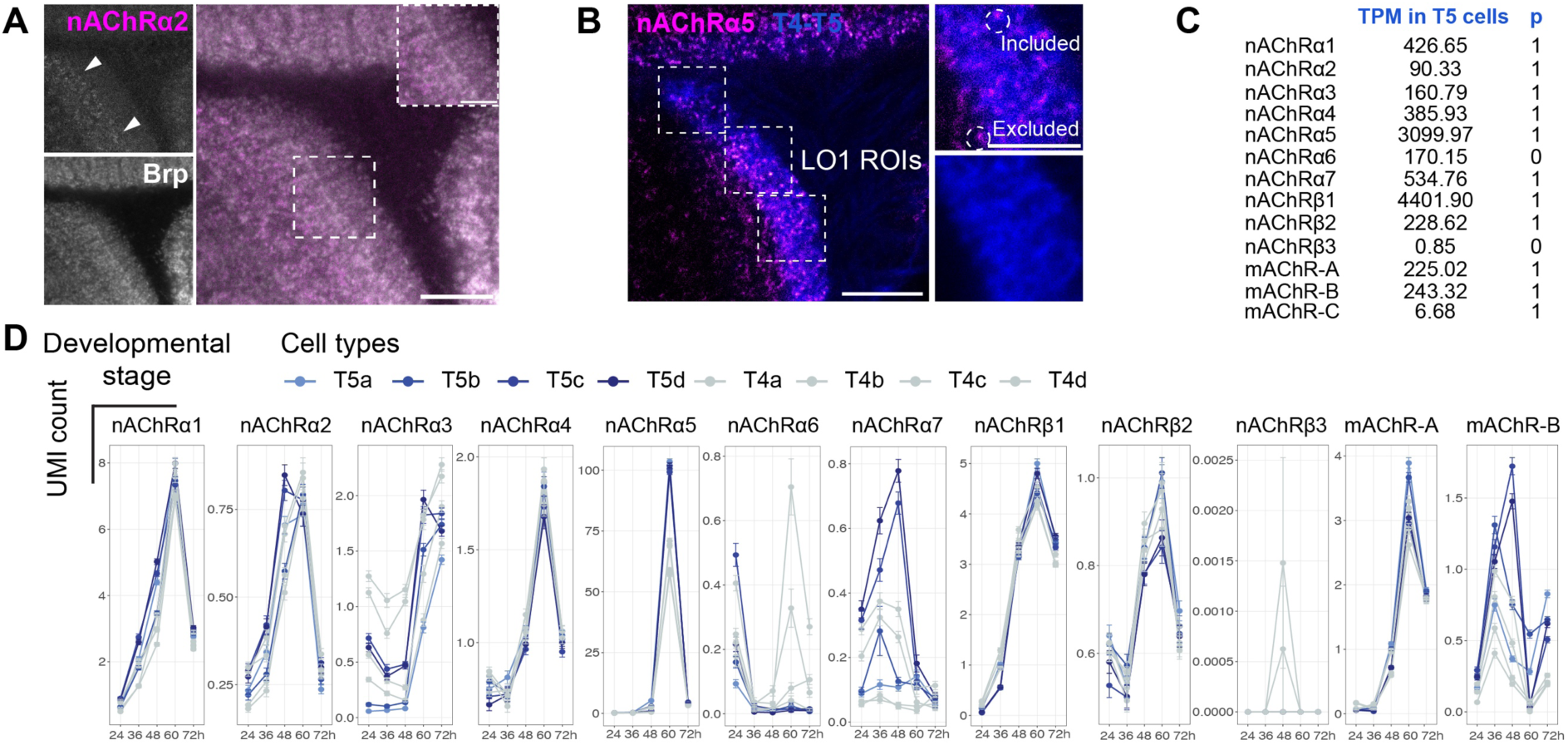
AChR expression in T5 cells-Related to Figure 1. (A) nAChRα2 expression in LO. Arrows indicate the LO layers of expression. Scale bar 10μm, inset 5μm. (B) Example ROIs in LO1 for quantification as in Figure 1J. Scale bar 10μm, inset 5μm. (C) Bulk RNAseq of AChR expression (transcripts per million, TPM) in T5 cells and probability of expression (p)^43^. (D) scRNAseq of AChR expression in T4 (grey) and T5 (blue) cells across five developmental stages (hours correspond to APF-after puparium formation)^58^.

**Figure S2.**
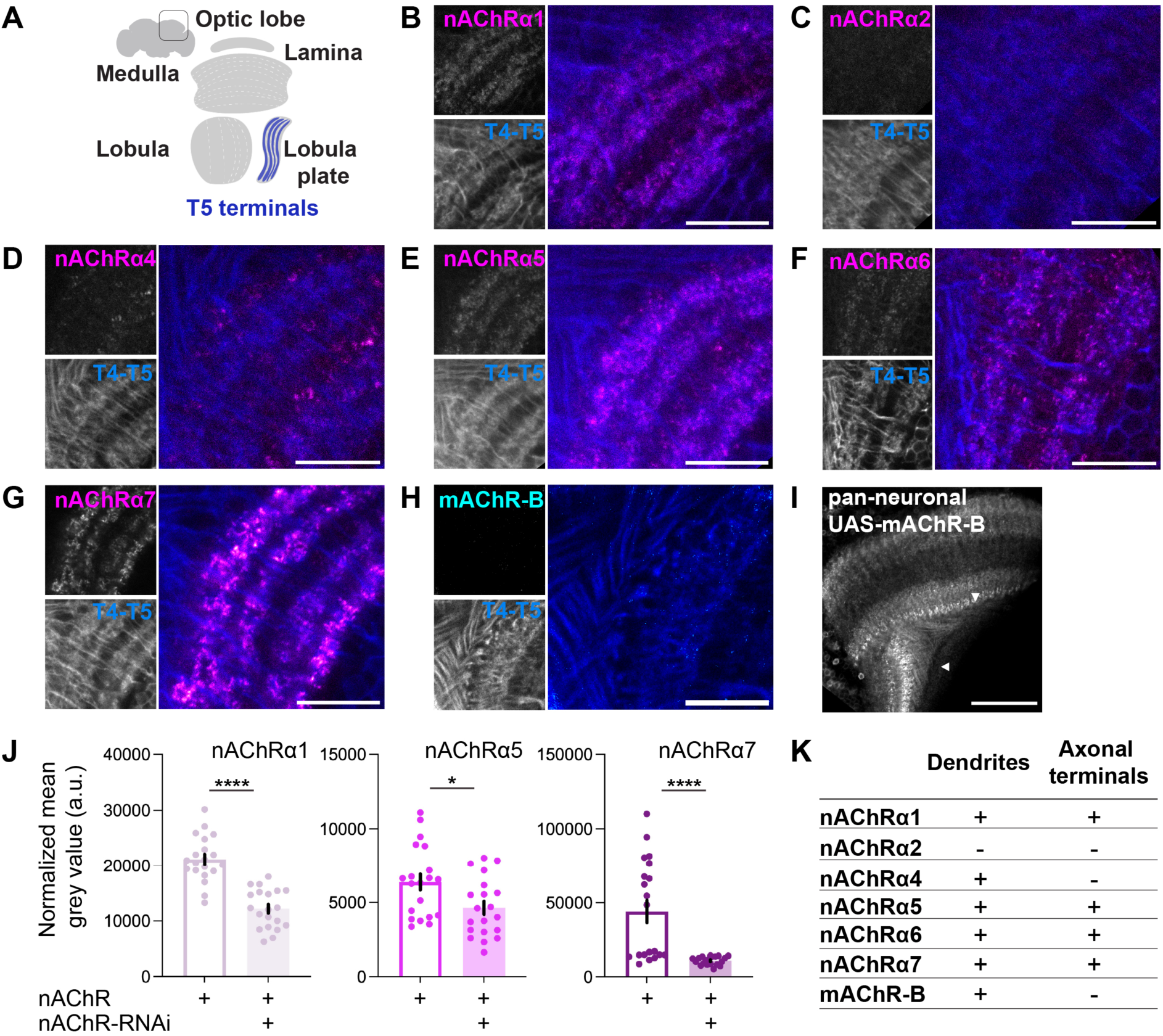
AChR expression in T4-T5 axonal terminals-Related to Figure 1. (A) Schematic representation of T5 axonal terminals residing in lobula plate (LOP1-4) of the *Drosophila* optic lobe (OL). (B-G) Expression of nAChRα1, nAChRα2, nAChRα4, nAChRα5, nAChRα6 and nAChRα7 (nAChRα::EGFP, magenta) in T4-T5 axonal terminals (R42F06-Gal4>UAS-myr::tdTomato, blue). Scale bar 10μm. (H) Overexpression of mAChR-B (cyan) in T4-T5 axonal terminals (R42F06-Gal4>UAS-myr::tdTomato, UAS-mAChR-B::HA). Scale bar 10μm. (I) Pan-neuronal expression of mAChR-B in OL (GMR57C10-Gal4>UAS-mAChR-B::HA). Scale bar 40μm. (J) Normalized mean grey value of nAChR subunits in LOP (layers 1 and 2) before and after RNAi induction (optic lobes n=2, ROIs N=20, for nAChRα1 n=1). The normality of distribution was assessed with the use of Shapiro-Wilk test. *p<0.05, ****p<0.0001. Data is mean ± SEM. (K) Summarized expression patterns of nAChRα1, nAChRα2, nAChRα4, nAChRα5, nAChRα6 and nAChRα7 subunits and the mAChR-B type on T5 dendrites and in T4-T5 axonal terminals.

**Figure S3.**
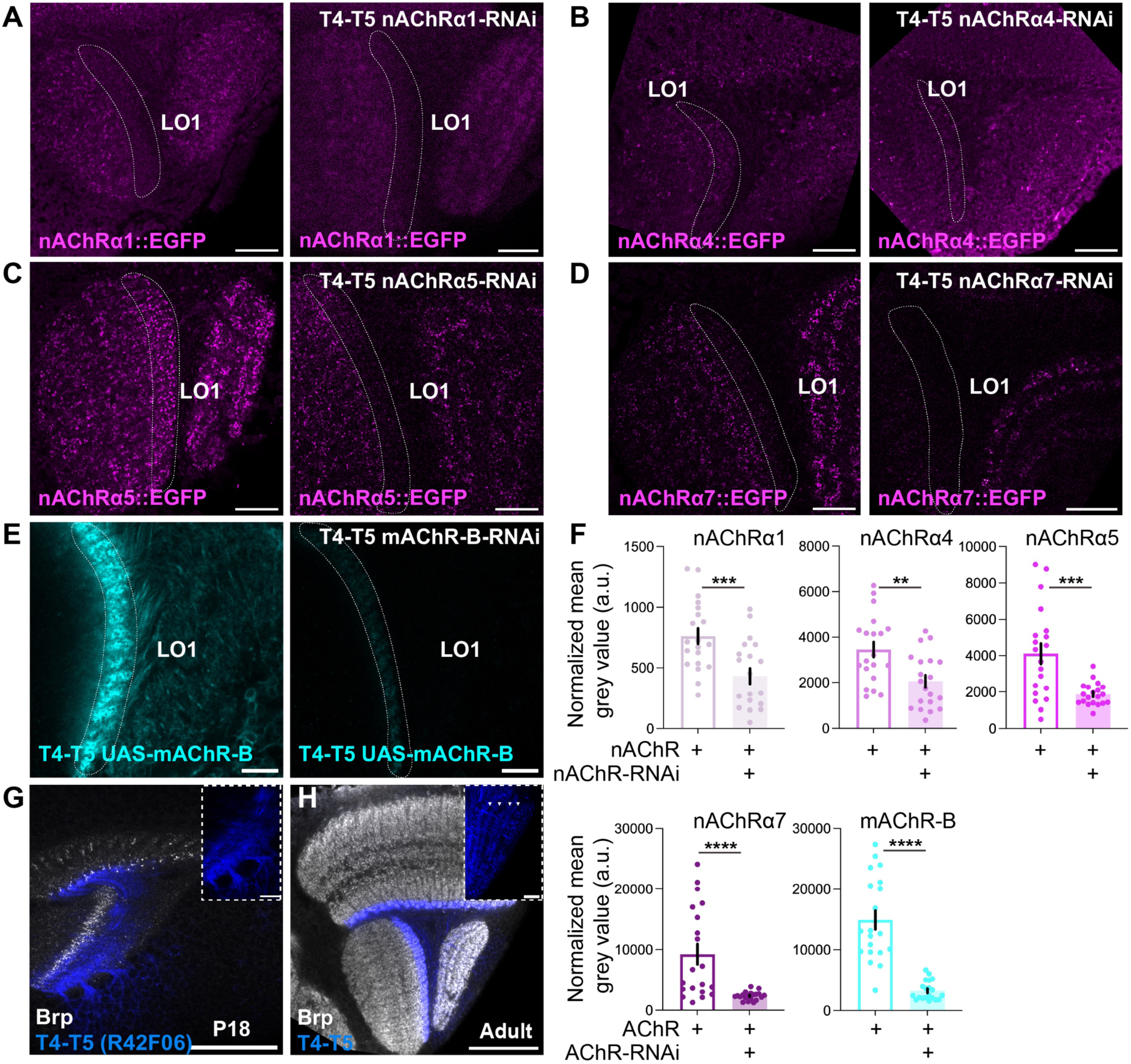
RNA interference against AChRs in T5 cells-Related to Figure 1 and 5. (A) Expression of the nAChRα1 subunit (nAChRα1::EGFP) in LO1 before (left panel) and after (right panel) the RNAi induction (R42F06-Gal4>AChR-RNAi). Scale bar 10μm. (B-E) As in (A) for the nAChRα4, nAChRα5, nAChRα7 subunits and the mAChR-B type. (F) Normalized mean grey value of AChRs in LO1 before and after RNAi induction (optic lobes n=2, ROIs N=20). The normality of distribution was assessed with the use of Shapiro-Wilk test. **p<0.01, ***p<0.001, ****p<0.0001. Data is mean ± SEM. (G) Developmental onset (P18) of the R42F06 promoter used for the split T4-T5 line in all behavioral experiments (R42F06-Gal4> UAS-myr::tdTomato). Scale bar 40μm, inset 10μm. (H) Split T4-T5 line (R42F06-p65.AD; VT043070-Gal4.DBD>UAS-CD4::tdGFP) used in all behavioral experiments. Arrows indicate the four LOP layers. Scale bar 40μm, inset 10μm.

**Figure S4.**
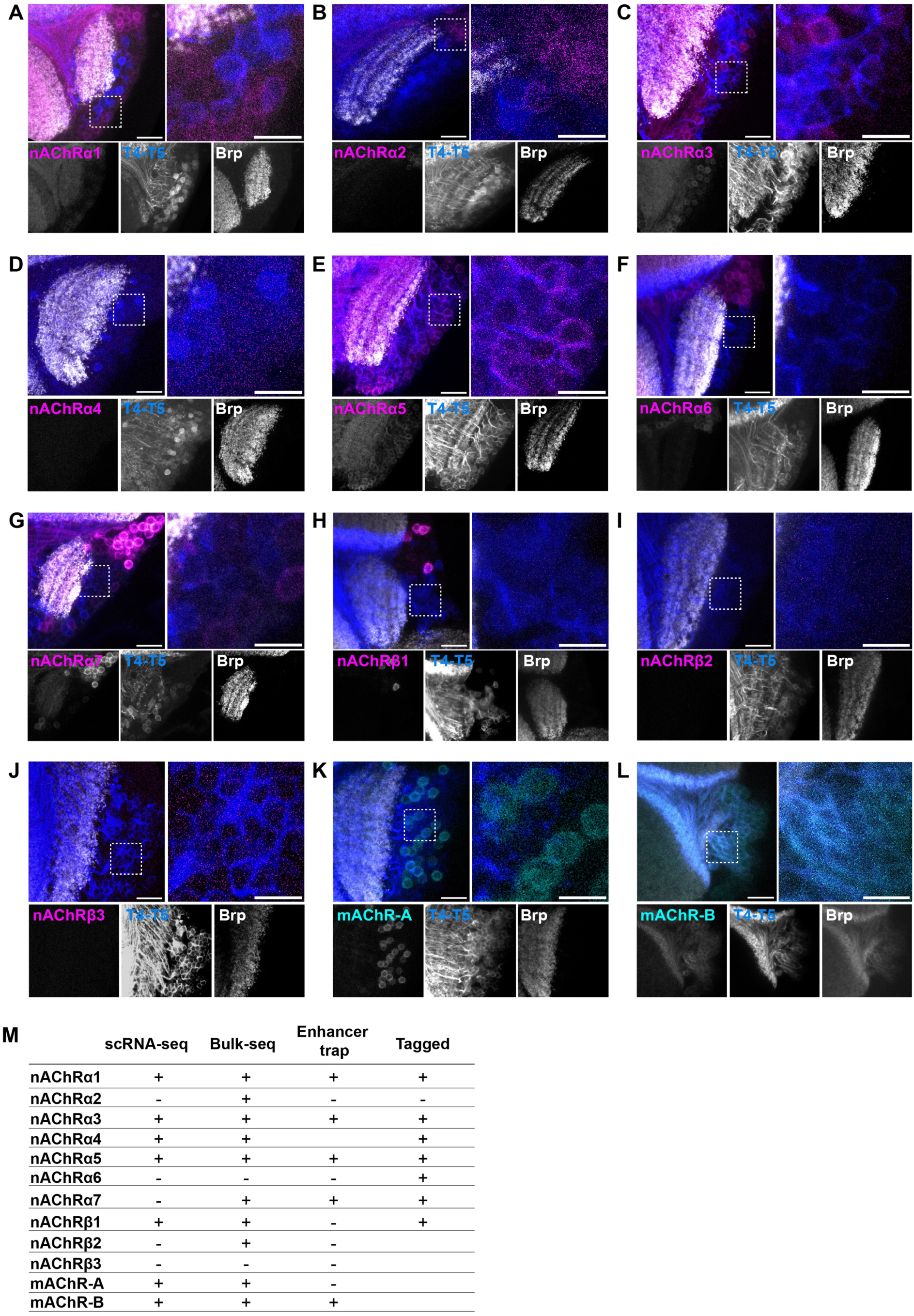
Enhancer-trap verification of AChR expression in T4-T5 cells-Related to Figure 1. (A-L) Enhancer-trap lines for the nAChRα1, nAChRα2, nAChRα3, nAChRα4, nAChRα5, nAChRα6, nAChRα7, nAChRβ1, nAChRβ2, nAChRβ3 subunits and mAChR-A and mAChR-B receptors (AChR-Mi or T2A-Gal4>UAS-CD8::GFP, magenta and cyan). Signal overlap with T4-T5 somata (R42F06-LexA>LexAop-CD8::RFP, blue) denoted the T4-T5 AChR expression. The nAChRα4 line did not show expression across all brain areas. Images are z-projections. Scale bar 40μm, inset 10μm. (M) AChR expression in T5 cells across scRNA-seq^58^, Bulk-seq^43^, enhancer trap and endogenously tagged receptor^60^ experiments.

**Figure S5.**
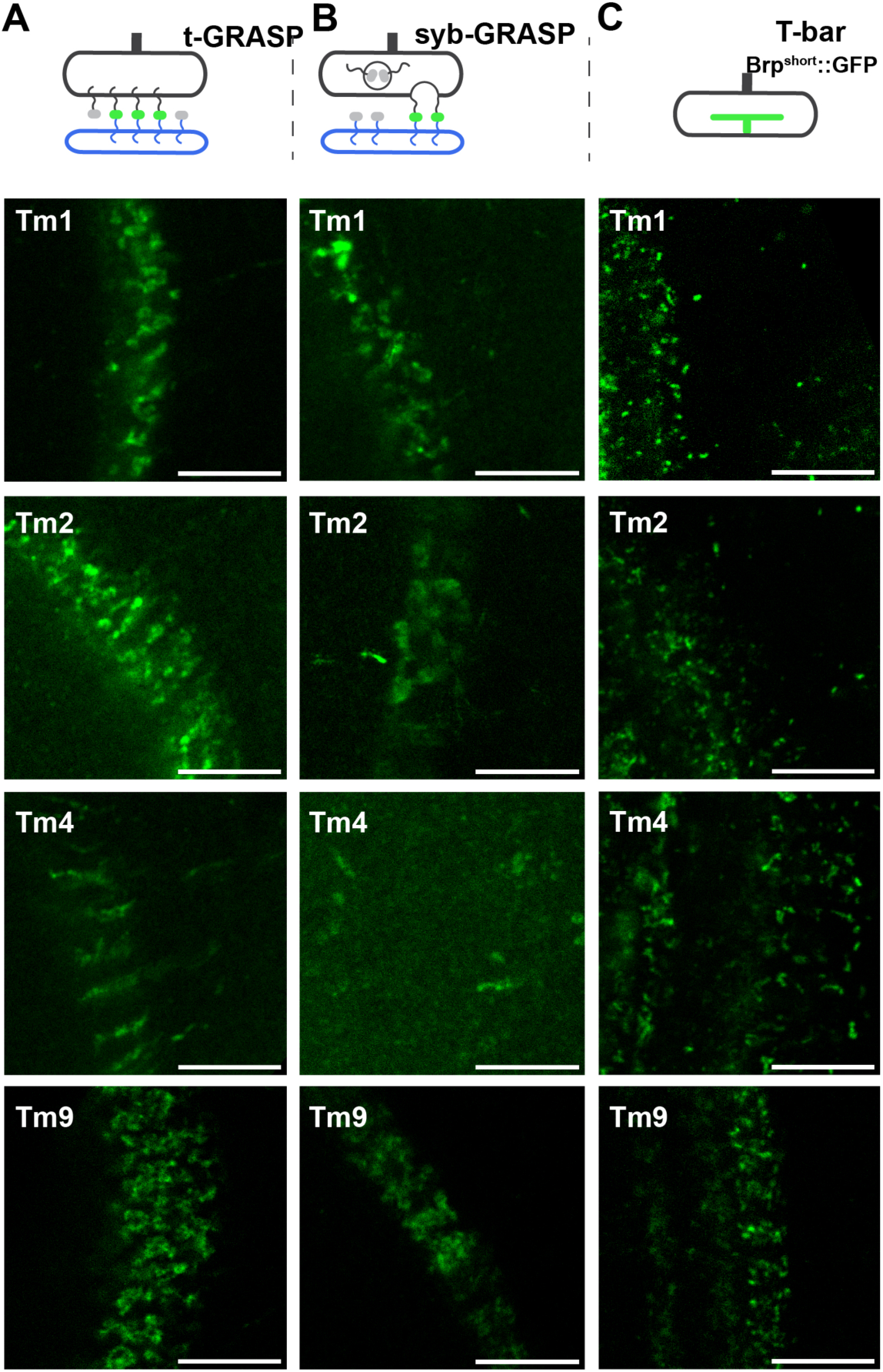
Screening of genetic approaches for synapse visualization-Related to Figure 3. (A) t-GRASP expression in Tm1, Tm2, Tm4 and Tm9-T5 synapses (Tm1, Tm2, Tm4, Tm9-Gal4>UAS-pre-t-GRASP, VT25965-LexAp65>LexAop-post-t-GRASP). Scale bar 10μm. Ectopic reconstitution is observed in Tm4-T5 synapses. (B) Activity GRASP (syb-GRASP) in Tm1, Tm2, Tm4 and Tm9-T5 synapses (Tm1, Tm2, Tm4, Tm9-Gal4>UAS-nSyb-spGFP1-10, VT25965-LexAp65>LexAop-CD4-spGFP11). Scale bar 10μm. (C) Brp^short^::GFP expression in Tm1, Tm2, Tm4 and Tm9 boutons (Tm1, Tm2, Tm4, Tm9-Gal4>UAS-Brp^short^::GFP). Scale bar 10μm.

**Figure S6.**
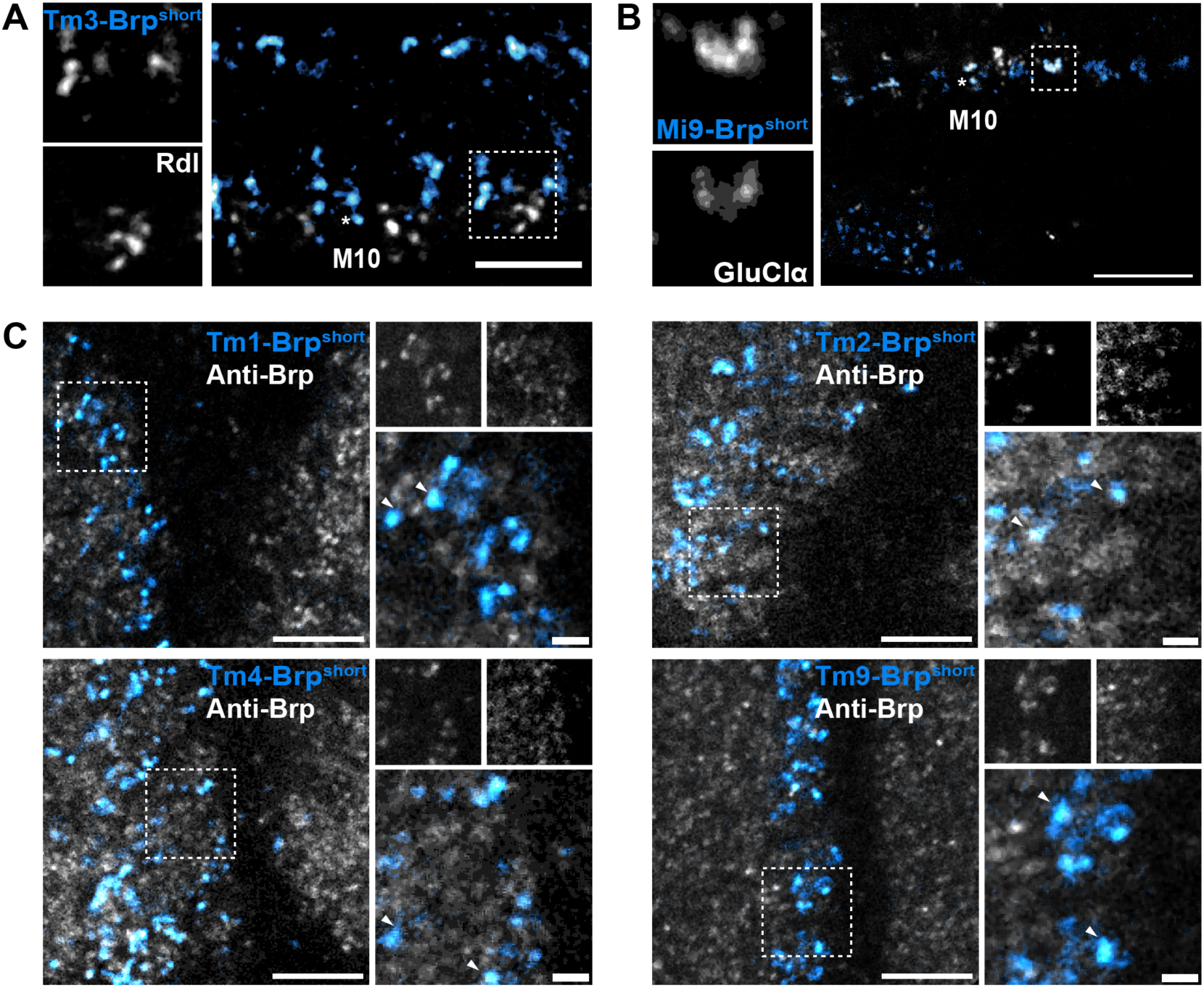
Validation of Brp^short^ as a precise presynaptic marker-Related to Figure 3. (A) Trans-synaptic alignment between Brp^short::mCherry^(blue) and the GABAergic Rdl receptor (grey) in the cholinergic Tm3-T4 synapse (GMR13E12-LexA>LexAop-Brp^short^::mCherry, R42F06-Gal4>UAS-Rdl::GFP). Scale bar 10μm. (B) Trans-synaptic alignment between Brp^short::mCherry^(blue) and the glutamatergic GluClα receptor (grey) in the glutamatergic Mi9-T4 synapse (GMR48A07-LexA>LexAop-Brp^short^::mCherry, R42F06-Gal4>UAS-GluClα::GFP). Scale bar 10μm. (C) Localization of Brp^short::mStraw^ in Tm1, Tm2, Tm4 and Tm9 boutons (blue) and endogenous Brp (grey) identified by the anti-Brp staining. Arrows correspond to the signal overlap between the Tm-Brp^short^ and the Anti-Brp. Scale bar 5 and 1μm.

**Figure S7.**
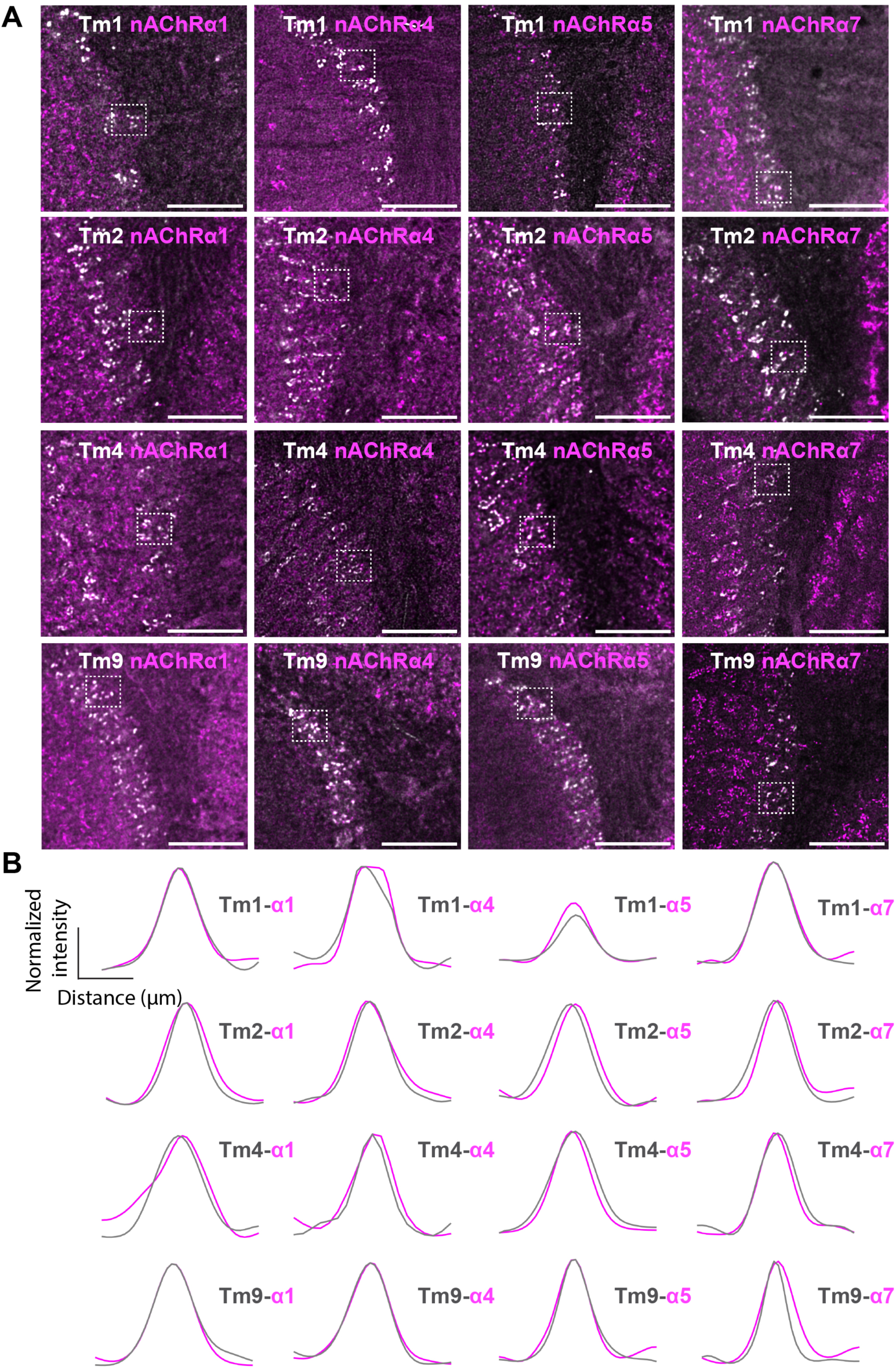
Localization of nAChRα subunits in Tm synapses-Related to Figure 3. (A) LO1 loci that were used for Figure 3A-D. Scale bar 10μm. (B) Line profile analysis for the assessment of trans-synaptic alignment of Brp^short::mStraw^ puncta (grey) and nAChRα puncta (magenta) across Tm1, Tm2, Tm4 and Tm9 synapses (analyzed puncta n=10). The dark lines represent the mean values.

**Figure S8.**
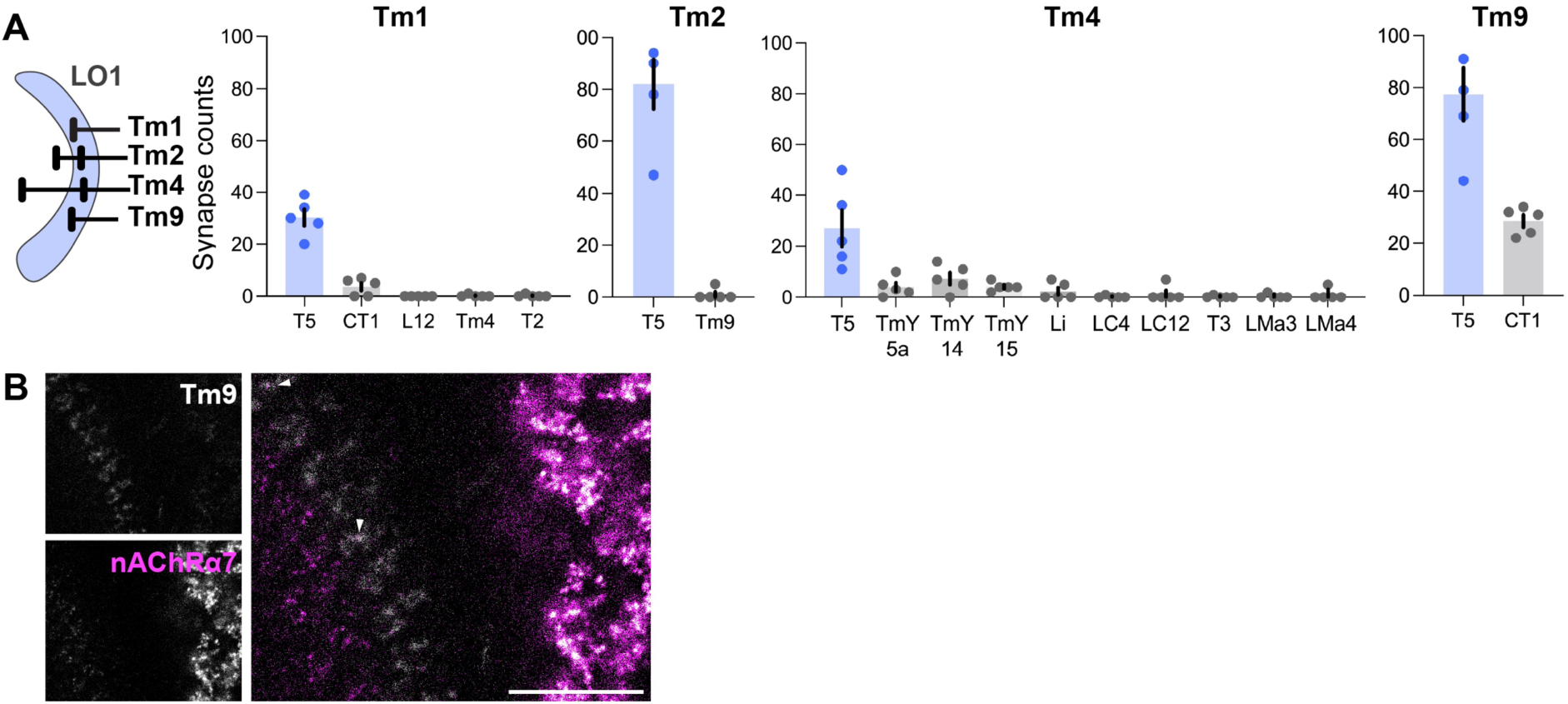
Neuronal outputs of Tm1, Tm2, Tm4 and Tm9 cells in LO1-Related to Figure 3. (A) Schematic representation of Tm1, Tm2, Tm4 and Tm9 axonal terminals in relation to LO1. Primary Tm1, Tm2, Tm4 and Tm9 output neuronal types in LO1 (Tm cells n=5, Codex). Data is mean ± SEM. (B) C-RASP for the Tm9-T5 synapse visualization (Tm9-Gal4>UAS-IVS-syb-spCFP1-10, VT25965-LexAp65>LexAop-CD4-spGFP11, grey) of the nAChRα7 subunit (nAChRα7::EGFP, magenta). Arrows correspond to Tm9-nAChRα7 trans-synaptic alignment. Scale bar 10μm.

**Figure S9.**
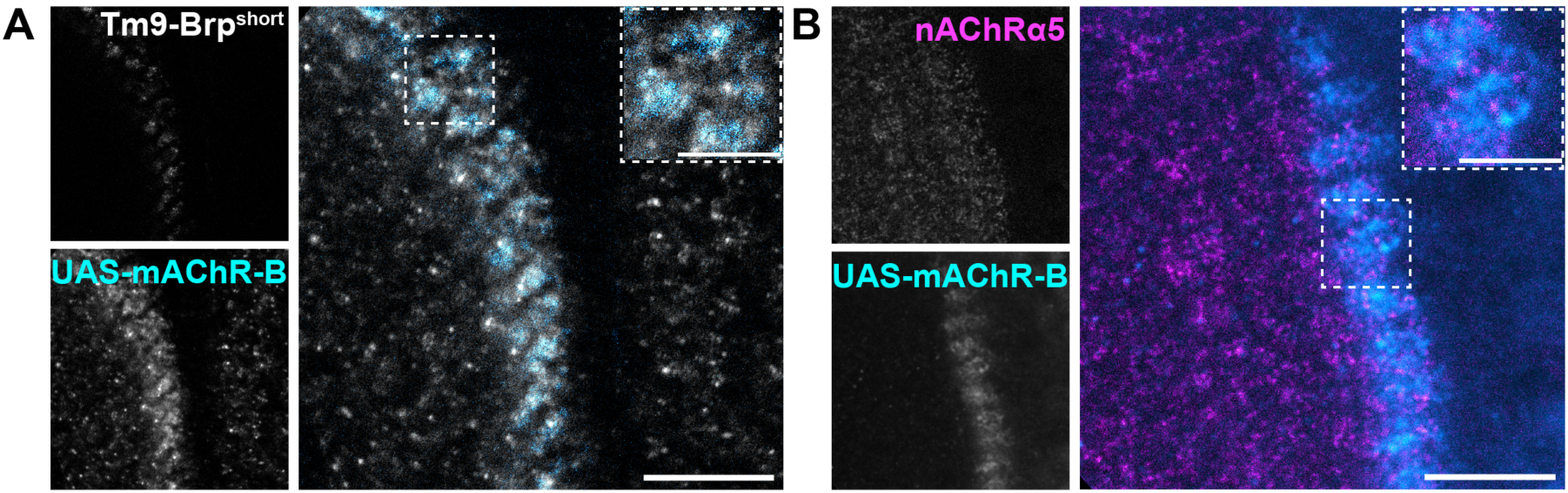
mAChR-B receptor localization in T5 dendrites-Related to Figure 3. (A) mAChR-B localization (R42F06-Gal4>UAS-mAChR-B::HA, cyan) in Tm9-T5 synapses (Tm9-LexA>LexAop-Brp^short^::mCherry, grey). Scale bar 10μm. (B) nAChRα5 subunit (nAChRα5::EGFP, magenta) localization pattern compared to mAChR-B (R42F06-Gal4>UAS-mAChR-B::HA, cyan) in LO1. Scale bar 10μm.

**Figure S10.**
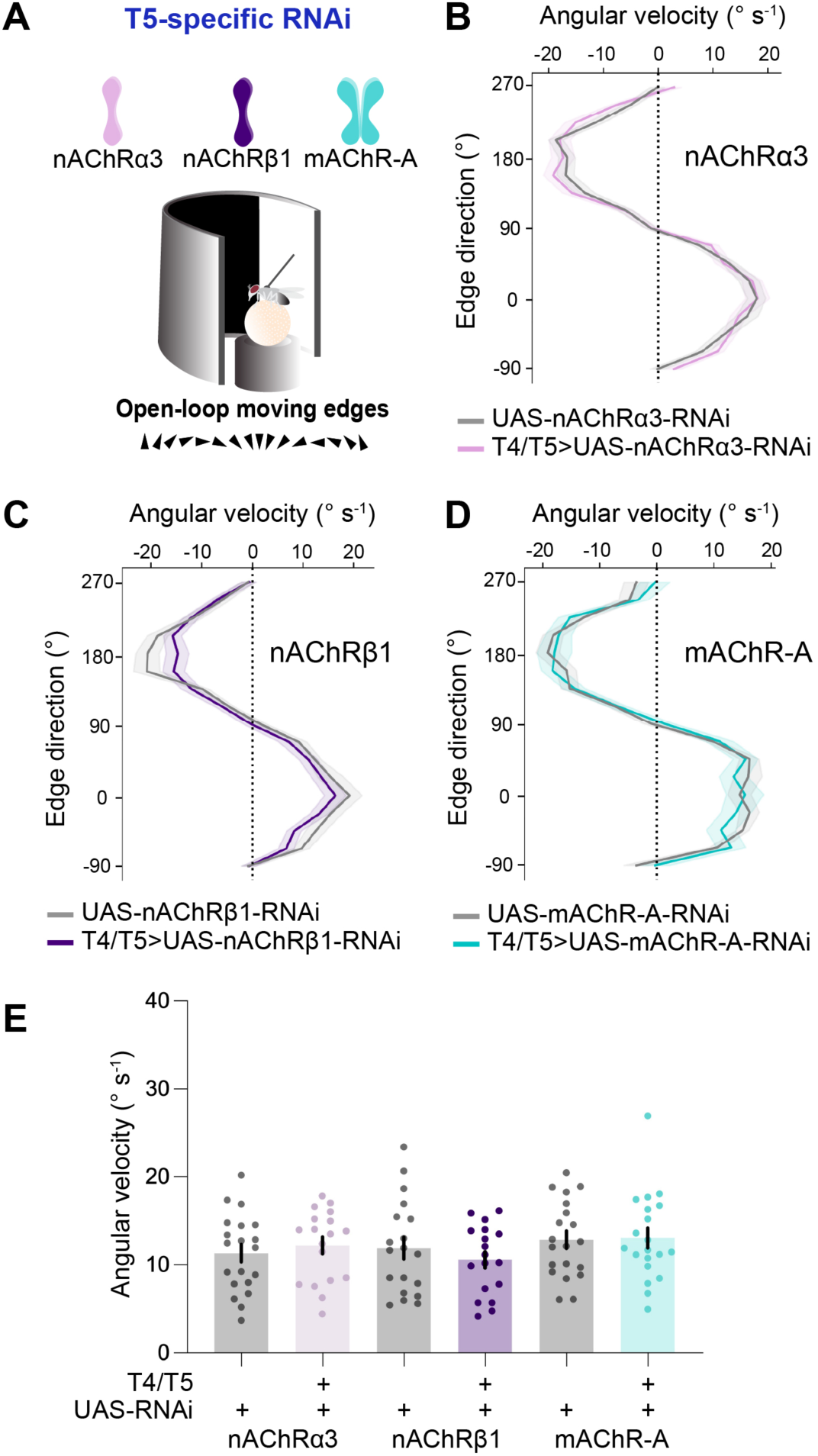
Fly optomotor response after nAChRα3, nAChRβ1 and mAChR-A knock-down-Related to Figure 5. (A) Schematic representation of the behavioral setup for the optomotor response assessment in nAChRα3 (pale magenta), nAChRβ1 (violet) subunits and mAChR-A (teal) type knock-down flies. (B) Trajectory (angular velocity) of nAChRα3 knock-down flies (42F06-p65.AD; VT043070-Gal4.DBD>UAS-nAChRα3-RNAi, pale magenta n=18) while presented to a sequence of upward to rightward to downward to leftward OFF edge motion compared to the parental control flies (UAS-nAChRα3-RNAi, grey n=20). The dark lines represent the mean ± SEM. (C-D) Same as previously for the nAChRβ1 (violet n=18, grey n=19) and mAChR-A (teal n=20, grey n=20) knock-down flies. (E) Angular velocity of control and AChR knock-down flies. The normality of distribution was assessed with the use of Shapiro-Wilk test. Data is mean ± SEM.

**Figure S11.**
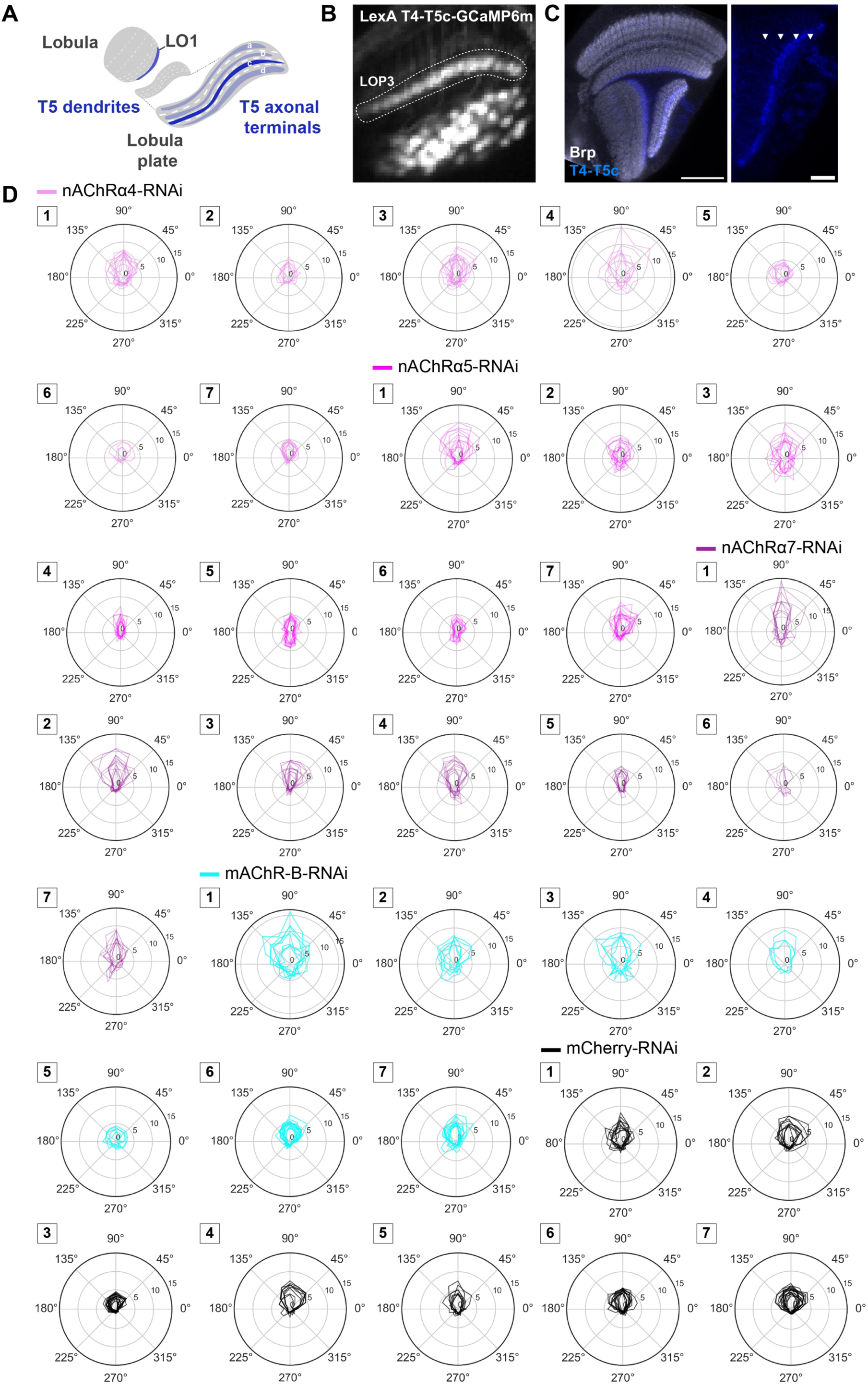
Two-photon calcium imaging in T5 cells-Related to Figure 6. (A) Schematic representation of LO1 (where T5 dendrites reside) and of LOP3 (where T5c axonal terminals reside). (B) Two-photon view of LOP3 (VT50384-LexA>LexAop-IVS-GCaMP6m). (C) T4-T5c specific targeting (VT50384-Gal4>UAS-CD8::GFP). Arrows indicate the four LOP layers. Scale bars 40 and 10μm. (D) Polar plots of the maximum responses (ΔF/F) -along seven flies-across sixteen OFF edge directions (°) of T5c cells under nAChRα4 RNAi (light magenta), nAChRα5 RNAi (magenta), nAChRα7 RNAi (dark magenta), mAChR-B RNAi (cyan) and control mCherry RNAi (black) conditions.

**Figure S12.**
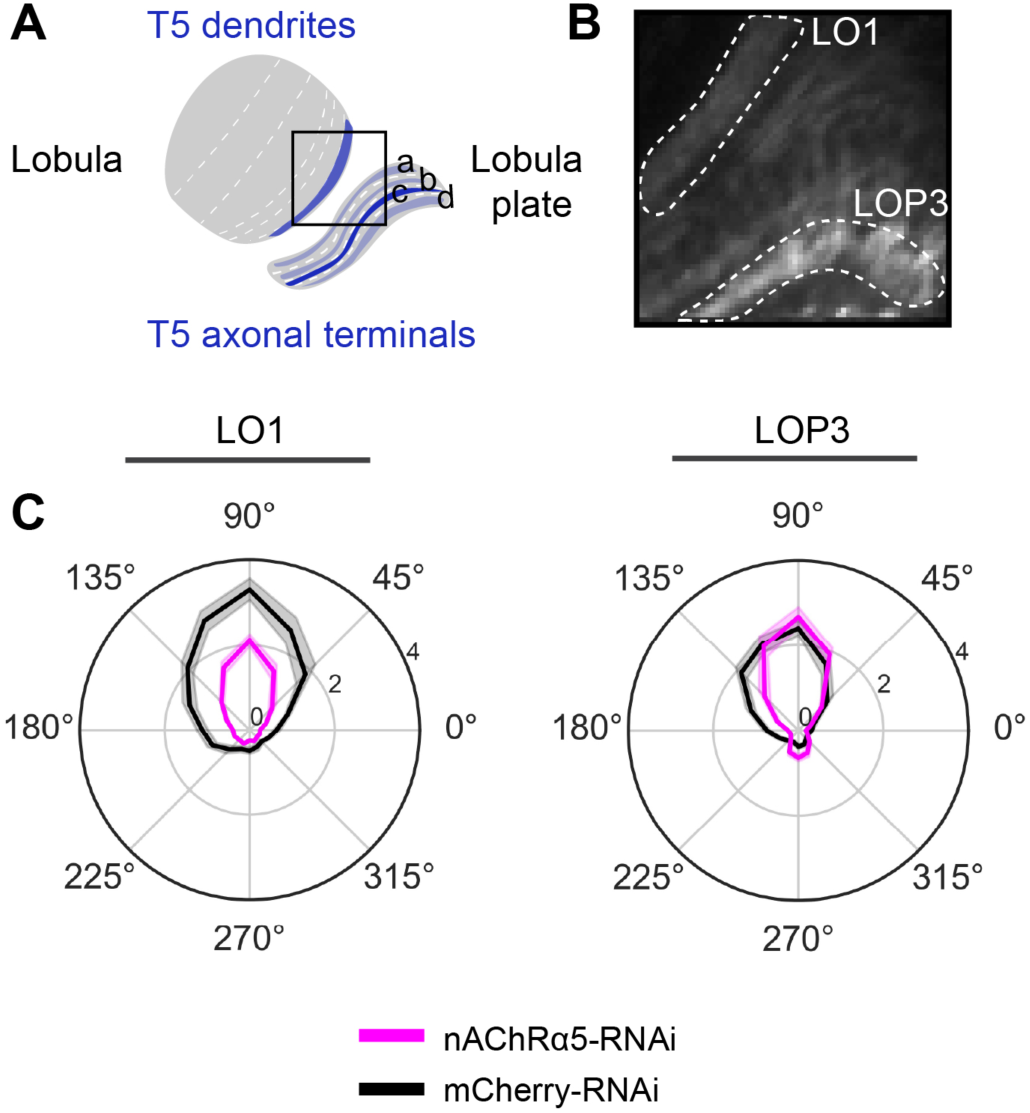
Two-photon calcium imaging in T5 dendrites and T5 axonal terminals-Related to Figure 6. (A) Schematic representation of LO1 (where T5 dendrites reside) and of LOP3 (where T5c axonal terminals reside). (B) Two-photon view of the LO1 and LOP3 layers (VT50384-LexA>LexAop-IVS-GCaMP6m). (C) Polar plots of the maximum responses (ΔF/F) across sixteen OFF edge directions (°) of T5c cells under mCherry controls (black, flies n=5, ROIs N=10) and nAChRα5 RNAi (magenta, n=5, N=13) conditions in LO1 (left) and in LOP3 (right). The dark lines represent the mean (68% CI).

**Table S1. List of fly experimental genotypes, related to STAR Methods.**

**Table S2. Statistical analysis, related to STAR Methods.**

**Table S3. FlyWire neuronal IDs, related to STAR Methods.**

## Notes

### Competing Interest Statement

The authors have declared no competing interest.

### Summary of Updates

Revision of the complete manuscript text including main and supplementary figures.

